# IgDesign: *In vitro* validated antibody design against multiple therapeutic antigens using inverse folding

**DOI:** 10.1101/2023.12.08.570889

**Authors:** Amir Shanehsazzadeh, Julian Alverio, George Kasun, Simon Levine, Ido Calman, Jibran A. Khan, Chelsea Chung, Nicolas Diaz, Breanna K. Luton, Ysis Tarter, Cailen McCloskey, Katherine B. Bateman, Hayley Carter, Dalton Chapman, Rebecca Consbruck, Alec Jaeger, Christa Kohnert, Gaelin Kopec-Belliveau, John M. Sutton, Zheyuan Guo, Gustavo Canales, Kai Ejan, Emily Marsh, Alyssa Ruelos, Rylee Ripley, Brooke Stoddard, Rodante Caguiat, Kyra Chapman, Matthew Saunders, Jared Sharp, Douglas Ganini da Silva, Audree Feltner, Jake Ripley, Megan E. Bryant, Danni Castillo, Joshua Meier, Christian M. Stegmann, Katherine Moran, Christine Lemke, Shaheed Abdulhaqq, Lillian R. Klug, Sharrol Bachas

## Abstract

Deep learning approaches have demonstrated the ability to design protein sequences given backbone structures [1, 2, 3, 4, 5]. While these approaches have been applied *in silico* to designing antibody complementarity-determining regions (CDRs), they have yet to be validated *in vitro* for designing antibody binders, which is the true measure of success for antibody design. Here we describe *IgDesign*, a deep learning method for antibody CDR design, and demonstrate its robustness with successful binder design for 8 therapeutic antigens. The model is tasked with designing heavy chain CDR3 (HCDR3) or all three heavy chain CDRs (HCDR123) using native backbone structures of antibody-antigen complexes, along with the antigen and antibody framework (FWR) sequences as context. For each of the 8 antigens, we design 100 HCDR3s and 100 HCDR123s, scaffold them into the native antibody’s variable region, and screen them for binding against the antigen using surface plasmon resonance (SPR). As a baseline, we screen 100 HCDR3s taken from the model’s training set and paired with the native HCDR1 and HCDR2. We observe that both HCDR3 design and HCDR123 design outperform this HCDR3-only baseline. IgDesign is the first experimentally validated antibody inverse folding model. It can design antibody binders to multiple therapeutic antigens with high success rates and, in some cases, improved affinities over clinically validated reference antibodies. Antibody inverse folding has applications to both *de novo* antibody design and lead optimization, making IgDesign a valuable tool for accelerating drug development and enabling therapeutic design. The data generated in this study serve as a useful benchmark of diverse antibody-antigen interactions. We use this data to benchmark self-consistency RMSD (scRMSD), using ABodyBuilder2 [6], ABodyBuilder3 [7], and ESMFold [8], as a metric for assessing binding. We open source the code for IgDesign and the SPR datasets.

Code and datasets can be found at https://github.com/AbSciBio/igdesign.

## 1 Introduction

Protein inverse folding is the problem of predicting sequences that fold into a structure of interest. Recently, several generative models for this task have been proposed [1, 2, 3, 4, 5]. These models have broad applications to the protein engineering space. In particular, ProteinMPNN [1] has been successfully applied to applications including *de novo* binder design [9], *de novo* luciferase design [10], and design of soluble analogs of membrane proteins [11]. Multiple other protein inverse folding models have been introduced, including ESM-IF1 [2] which uses a GVP-GNN architecture [12] and LM-Design [3] combines structure encoders (such as ProteinMPNN) with the ESM protein language models [13, 14].

Antibodies have become a major class of therapeutics as a result of their attractive drug-like properties [15]. Antibody design, in particular design of the hypervariable CDRs primarily involved in antigen binding [16], using generative models is of great interest due to potential drug development applications. Correspondingly, antibody inverse folding has become a relevant problem. AbMPNN [5] describes an antibody-specific inverse folding model created by fine-tuning Protein-MPNN on antibody structures, such as those in the Structural Antibody Database (SAbDab) [17]. AntiFold [18] describes another antibody-specific inverse folding model created by fine-tuning ESM-IF1 on antibody structures. AbMPNN and AntiFold have been compared compared *in silico* to ProteinMPNN and ESM-IF1 and shown to have higher amino acid recovery (AAR) on antibody CDRs. AntiFold’s losses were also shown to be more predictive of antibody-antigen binding affinity compared to other models, however neither of the two models have been shown to produce antibody binders *in vitro*. Shanker et al. [19] affinity matured antibodies against SARS-CoV-2 with ESM-IF1 by ranking mutants using backbone structures, but they did not demonstrate the ability to design entire CDRs.

Our study introduces IgDesign, a generative antibody inverse folding model based on LM-Design, and the first such model to be validated *in vitro* for antibody binder design. We present extensive wet lab validation of IgDesign’s ability to design binders against 8 therapeutic antigens. Specifically, we show that IgDesign is able to design both HCDR3 and all three HCDRs (HCDR123) of a reference antibody^2^ and preserve binding, as measured by surface plasmon resonance (SPR), to the target antigen with high success rates. We compare IgDesign to a baseline of HCDR3s sampled from the model’s training set and observe that for HCDR3 design, the model outperforms on 8 out of 8 antigens. For HCDR123 design, the model outperforms this HCDR3-only baseline on 7 out of 8 antigens. Overall, we demonstrate, with *in vitro* validation, that inverse folding can be a successful component of an antibody design pipeline (Figure 1).

**Figure 1:**
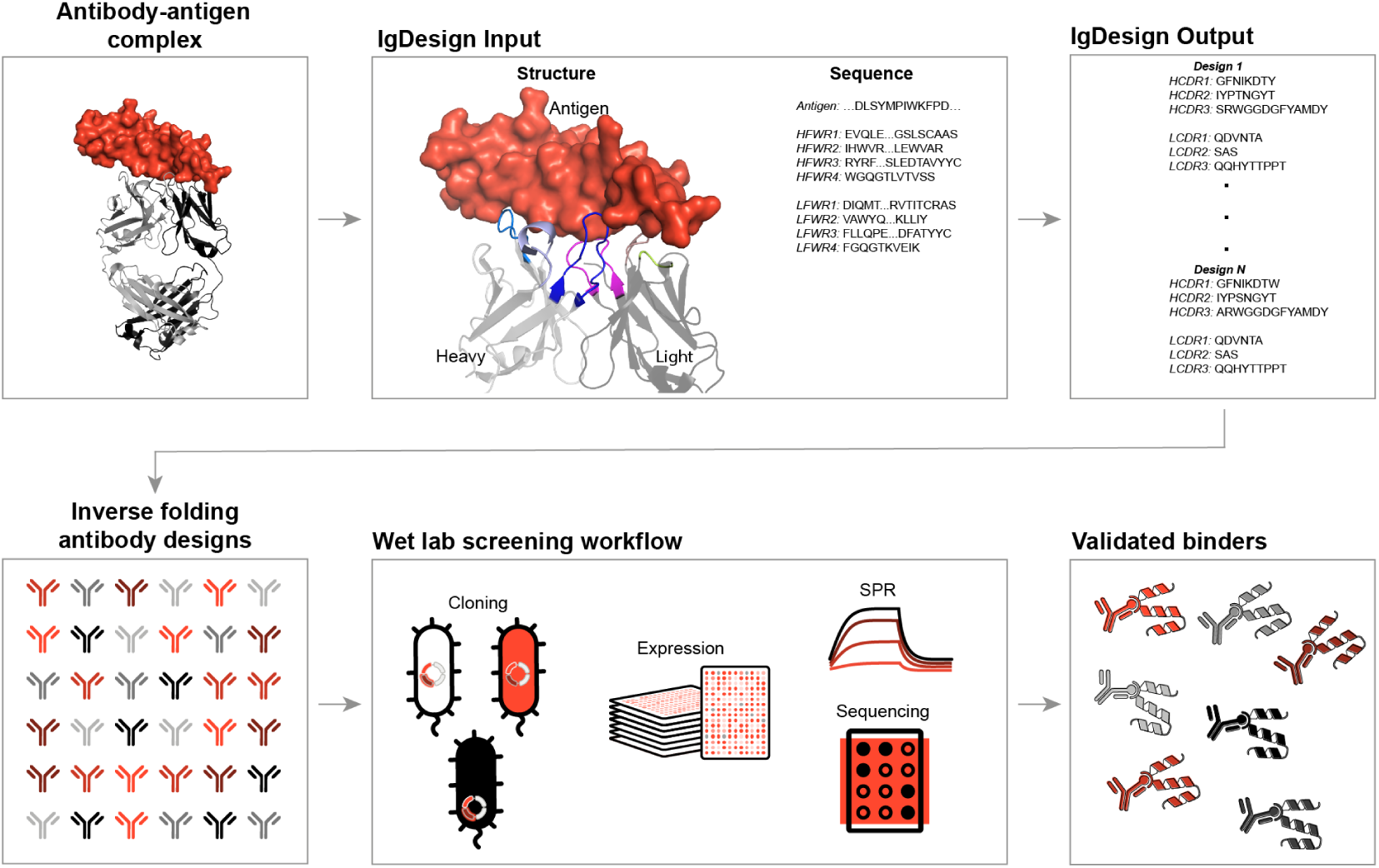
Overview of *in silico* (top) and *in vitro* (bottom) workflow for antibody inverse folding. (Top) Antibody-antigen complex crystal structures are processed into structure features (antibody variable-region and antigen coordinates) and sequence features (antigen and antibody framework sequences). Features are inputted to IgDesign which outputs CDR sequences. CDR loops are colored in the processed structure (PDB:1N8Z [20]). **(Bottom)** Libraries of inverse folding antibody designs are sent to the wet lab for screening. The screening workflow consists of cloning, expression, surface plasmon resonance (SPR), and sequencing. Designed binders are validated and their affinities are measured.

In evaluating IgDesign’s ability to design antibody binders, we, across 8 targets, screened 1437 antibodies via SPR and identified 278 binders. These datasets may be of great use in benchmarking antibody design models. We showcase this by evaluating the ability of the popular self-consistency RMSD (scRMSD) metric, computed using different folding models, to discriminate between binders and non-binders. We also measure correlation with affinity. We open source these datasets to facilitate future benchmarking efforts by the community.^3^

## 2 Methods

### IgDesign Model

To understand whether the success of ProteinMPNN [1] could be used in the antibody design field, we devised *IgMPNN*, a version specifically trained for antibodies. IgMPNN is similar to AbMPNN [5], however we note two key differences between the models: (1) IgMPNN is provided antigen sequence and antibody framework (FWR) sequences as context during training. (2) IgMPNN decodes antibody CDRs in sequential order during training: HCDR1, HCDR2, HCDR3, LCDR1, LCDR2, LCDR3. During inference, any order of CDRs can be specified. Within each CDR, the decoding order is random as done in [1]. We include additional details on IgMPNN in the Appendix.

The CDR design protocol in *IgDesign* is based on the approach of combining a structure encoder and sequence decoder as proposed in LM-Design [3]. We first execute a forward pass through IgMPNN, as described above. This allows us to access the final node embeddings as well as the model’s logits. We sample the maximum likelihood estimate of those logits in order to obtain a single tokenized sequence. We provide this sequence as input to the ESM2-3B protein language model [14]^4^ and extract the embeddings before the final projection head. We then apply a BottleNeck Adapter layer [21], in which cross-attention is computed by using the final node embeddings from IgMPNN as keys and the embeddings from ESM2-3B as queries and values. This new set of embeddings is passed through the final projection head of ESM2-3B and projected out to logits. Finally, these logits are summed with the logits from IgMPNN.

### Data and training

IgMPNN is pretrained on proteins from the PDB [22] split at 40% sequence identity (referred to here as the General PDB dataset). Following pretraining, IgMPNN is fine-tuned on antibody-antigen complexes from SAbDab [17] and IgDesign is fine-tuned on said antibody-antigen complexes using IgMPNN (pretrained and fine-tuned) as its structural encoder. For splitting the antibody data we do an antigen holdout at 40% sequence identity. We remove all structures in SAbDab from our General PDB dataset to avoid data leakage. We include additional details on data curation and splitting in the Appendix.

For each antigen, we train a new set of IgMPNN and IgDesign models with said antigen held out to prevent data leakage. We ensure that the HCDR3 of the reference antibody is not in the training set (Table 1). All models are trained with the Adam optimizer [23] using a learning rate of 10^−3^.

**Table 1:** Minimum edit distances between HCDRs of reference antibodies and antigen data splits used for training. Note that the reference antibody’s HCDR3 is never present in the training set. While HCDR1 and HCDR2 may be contained in the training set this is expected due to these being lower diversity regions relative to HCDR3. Furthermore, we note that the minimum edit distance between all three HCDRs is always greater than the sum of the minimum edit distances for each HCDR.

| Antigen | Minimum Edit Distance |  |  |  | Mean Edit Distance |  |  |  |
| --- | --- | --- | --- | --- | --- | --- | --- | --- |
|  | HCDR1 | HCDR2 | HCDR3 | HCDRs | HCDR1 | HCDR2 | HCDR3 | HCDRs |
| CD40 | 0 | 2 | 2 | 9 | 4.2 | 5.6 | 10.8 | 20.5 |
| IL36R | 1 | 2 | 7 | 13 | 5.2 | 5.7 | 12.7 | 23.6 |
| C5 | 1 | 0 | 5 | 10 | 4.7 | 4.8 | 12.1 | 21.6 |
| TSLP | 1 | 0 | 6 | 9 | 4.7 | 5.7 | 13.0 | 23.4 |
| IL17A | 0 | 2 | 8 | 13 | 4.5 | 5.6 | 13.3 | 23.4 |
| FXI | 1 | 1 | 4 | 11 | 4.4 | 4.6 | 11.9 | 20.9 |
| ACVR2B | 1 | 2 | 2 | 6 | 4.6 | 4.8 | 10.7 | 20.1 |
| TNFRSF9 | 0 | 0 | 2 | 7 | 4.7 | 5.3 | 10.5 | 20.5 |

### Antibody library design

We selected 8 therapeutic antigens each with a reference antibody binder and an antibody-antigen complex structure in our curated SAbDab dataset. The 3D backbone coordinates were used as input to IgDesign, along with the antigen sequence and antibody FWR sequences. At inference time, we generated sequences in the following order: HCDR3, HCDR1, HCDR2, LCDR1, LCDR2, LCDR3. For each antigen, we generated 1 million sequences and filtered to the 100 HCDR3s and 100 HCDR123s with lowest cross-entropy loss for *in vitro* assessment.

As a baseline, we sampled 100 HCDR3s from the training set (a subset of SAbDab) of each IgDesign model. SAbDab HCDR3s represent a rigorous baseline since they match the training distribution of the model and are paired with the native HCDR1 and HCDR2. In total, we designed 8 libraries of antibodies, one for each target antigen. Each antigen specific library includes 100 IgDesign HCDR3s, 100 IgDesign HCDR123s, and 100 SAbDab HCDR3s as a baseline. For each library, we included controls to confirm the SPR assay’s ability to accurately label known binders and non-binders. Additional details, including a discussion of the baseline, are in the Appendix.

### *In vitro* screening

Antibodies were screened for binding against their target antigens using SPR. We give a detailed overview of our wet lab workflow in the Appendix. At a high level, this workflow involves four steps: (1) *Cloning* DNA corresponding to the designed antibodies into *E. coli* plasmids (2) *Expressing* the antibodies in the corresponding *E. coli* (3) *SPR*: screening the expressed antibodies for binding against their target antigens (4) *Sequencing* the antibodies to determine amino acid identities. After receiving the SPR and sequencing data for each library, we label antibody variants that bind in all SPR replicates as binders and otherwise label them as non-binders.

## 3 Results

### Amino acid recovery (AAR)

We compute AAR, the percentage of the designed sequence that matches the native sequence, for each CDR using IgMPNN, IgDesign, and ProteinMPNN [1] as an *in silico* baseline^5^ We also evaluate “ProteinMPNN (Filtered),” which is ProteinMPNN evaluated on antibodies not in its training set. We compute 1-shot AAR, the AAR of one sample from the model. For each of the 8 antigens, we trained a model with said antigen held out and computed test set AARs. To estimate model performances, we compare mean test set AARs, that is the mean computed over each model’s test set. We show the distribution of mean test set 1-shot AARs for HCDRs in Figure 3 and for LCDRs in Figure 4. IgMPNN and IgDesign outperform ProteinMPNN and ProteinMPNN (Filtered) in all cases (Mann-Whitney U test (MWU) [24], *p* < 2e-4). IgDesign outperforms IgMPNN (MWU, *p* < 7e-4) on LCDR3. We compare test set AARs on the ACVR2B data split in Figures 5, 6.

**Figure 2:**
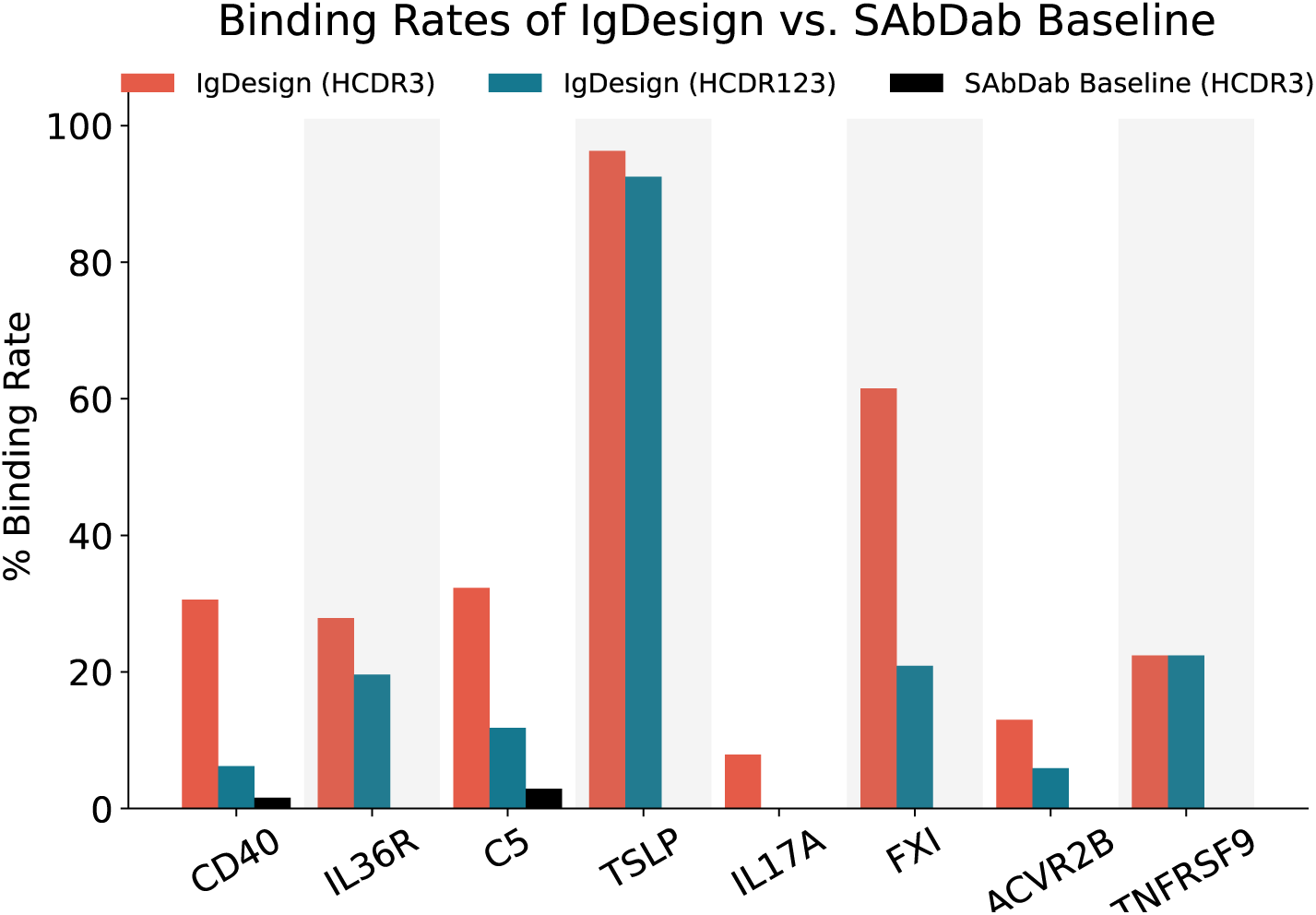
Binding rates across antigens for IgDesign HCDR3 (red) and HCDR123 (blue) vs. SAbDab HCDR3 baseline (black). Binding rate is defined as the percentage of sequences that bind to the target antigen as assessed by SPR. Baseline binding rate is 0% for IL36R, TSLP, IL17A, FXI, ACVR2B, and TNFRSF9. IgDesign significantly outperforms the SAbDab baseline.

**Figure 3:**
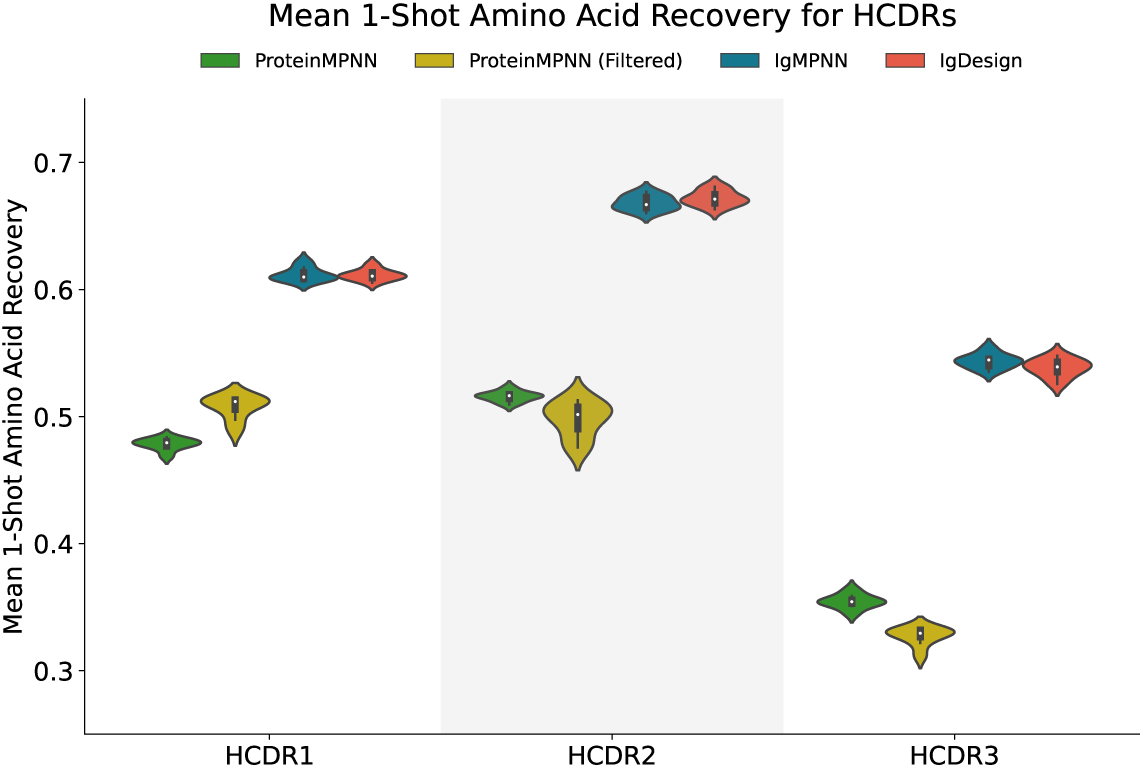
Comparison between IgMPNN and IgDesign on mean 1-shot amino acid recovery (AAR) for heavy chain CDRs (HCDRs). Violin plots comparing distributions of mean 1-shot AARs for ProteinMPNN (green), ProteinMPNN filtered to complexes not in its training set (yellow), IgMPNN (blue), and IgDesign (red) for each HCDR across 8 antigen test sets. Mean 1-shot AAR is the mean of the 1-shot AARs computed across each test set. The distribution captures the 95% interval, the white dot represents the median, and the box represents the interquartile range.

**Figure 4:**
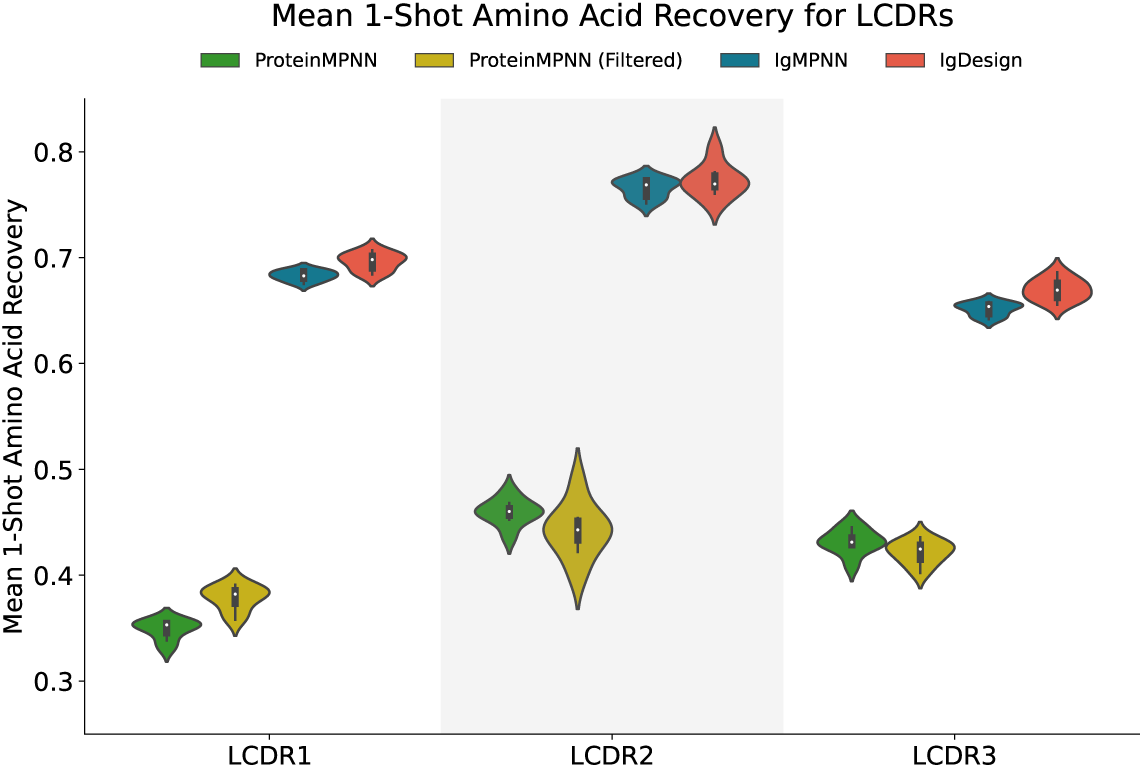
Comparison between IgMPNN and IgDesign on mean 1-shot amino acid recovery (AAR) for light chain CDRs (LCDRs). Violin plots comparing distributions of mean 1-shot AARs for ProteinMPNN (green), ProteinMPNN filtered to complexes not in its training set (yellow), IgMPNN (blue), and IgDesign (red) for each LCDR across 8 antigen test sets. Mean 1-shot AAR is the mean of the 1-shot AARs computed on each test set. The distribution captures the 95% interval, the white dot represents the median, and the box represents the interquartile range.

**Figure 5:**
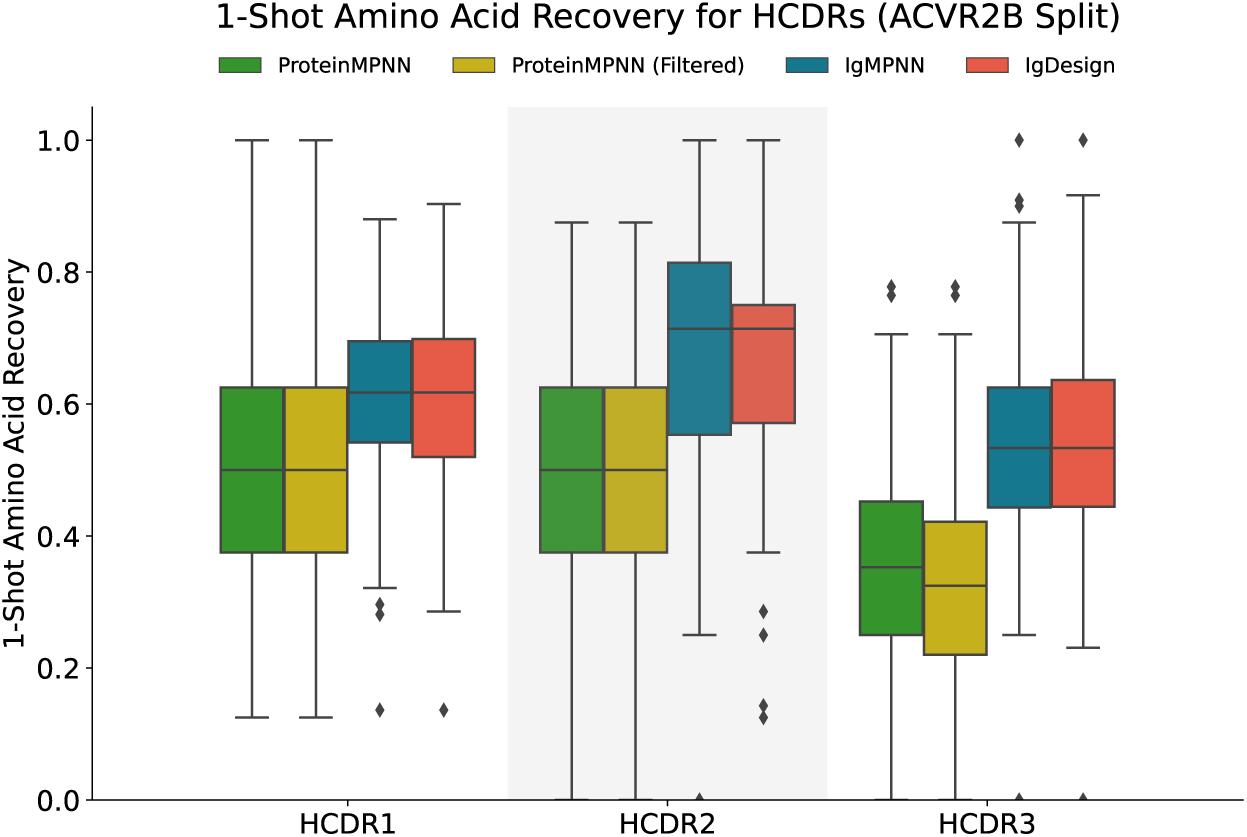
Comparison between IgMPNN and IgDesign on 1-shot amino acid recovery (AAR) for heavy chain CDRS (HCDRs) on ACVR2B test set. Box plots comparing distributions of 1-shot AARs for ProteinMPNN (green), ProteinMPNN filtered to complexes not contained in its training set (yellow), IgMPNN (blue) and IgDesign (red) across the test set (ACVR2B data split of SAbDab) for each HCDR.

**Figure 6:**
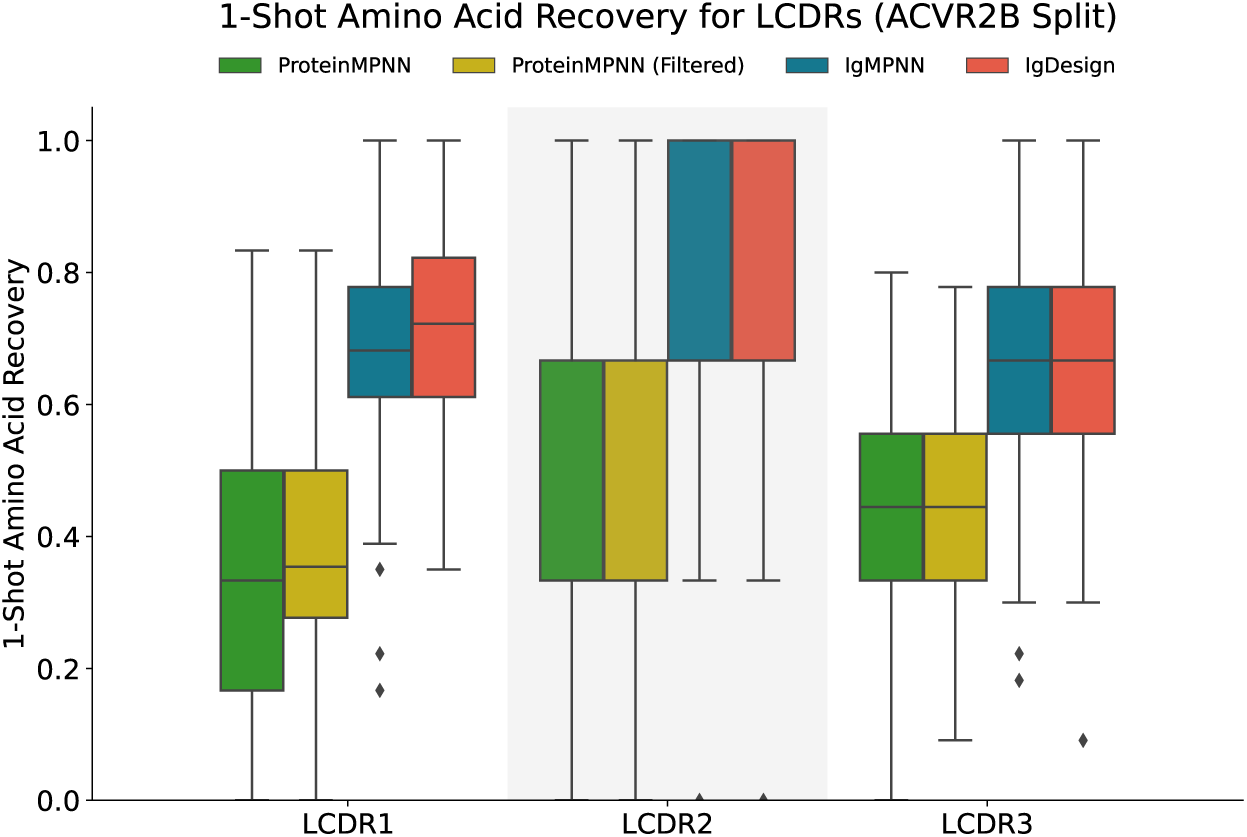
Comparison between IgMPNN and IgDesign on 1-shot amino acid recovery (AAR) for light chain CDRS (LCDRs) on ACVR2B test set. Box plots comparing distributions of 1-shot AARs for ProteinMPNN (green), ProteinMPNN filtered to complexes not contained in its training set (yellow), IgMPNN (blue) and IgDesign (red) across the test set (ACVR2B data split of SAbDab) for each LCDR.

### *In vitro* binding rates and affinities

We assess IgDesign’s ability to generate binders to each therapeutic antigen by measuring its *in vitro* binding rate, which is the percentage of designs that bind as assessed by SPR. As shown in Figure 2 and Tables 3, 4, IgDesign HCDR3s bind statistically significantly more often than SAbDab baseline HCDR3s for 7 out of 8 antigens (Fisher’s exact test (FE) [25], *p* < 3e-3)^6^. IgDesign HCDR123s bind statistically significantly more often than the baseline for 4 out of 8 antigens (FE, *p* < 3e-3). Baseline HCDR3s bound for 2 antigens (1 against CD40 and 2 against C5). We emphasize the impact of designing HCDR123s with high binding rates as we are comparing to an HCDR3-only baseline. IgDesign’s significant outperformance vs. the baseline suggests it has learned to extrapolate from its training set. For 5 out of 8 antigens (CD40, IL36R, FXI, ACVR2B, TNFRSF9), IgDesign generated binders with equal or higher affinities to the reference antibody. For the other 3 out of 8 antigens (C5, TSLP, IL17A), the binders are 2+ orders of magnitude lower affinity than reference, motivating future work to improve the model. We report and discuss binding affinities in Figures 7, 8, 9, 10, 11, 12, 13, 14 and in the Appendix.

**Table 2:** PDB IDs and chain IDs for antibody-antigen complexes.

| **Antigen** | **Reference Antibody** | **PDB ID** | **Heavy Chain ID** | **Light Chain ID** | **Antigen Chain ID** |
| --- | --- | --- | --- | --- | --- |
| CD40 | Ravagalimab | 6PE8 | A | B | U |
| IL36R | Spesolimab | 6U6U | H | L | R |
| C5 | Eculizumab | 5I5K | H | L | B |
| TSLP | Tezepelumab | 5J13 | C | B | A |
| IL17A | Afasevikumab | 6PPG | B | A | G |
| FXI | Osocimab | 6HHC | H | L | A |
| ACVR2B | Bimagrumab | 5NGV | H | L | A |
| TNFRSF9 | Utomilumab | 6A3W | A | B | C |

**Table 3:** Binding rates across antigens for IgDesign on HCDR3 and HCDR123 as well as SAbDab HCDR3 baseline.

| Antigen | % Binding Rate (Binders / Observations) |  |  |
| --- | --- | --- | --- |
|  | IgDesign (HCDR3) | IgDesign (HCDR123) | SAbDab (HCDR3) |
| CD40 | 30.6% (22 / 72) | 6.3% (4 / 64) | 1.6% (1 / 64) |
| IL36R | 27.9% (17 / 61) | 19.6% (11 / 56) | 0.0% (0 / 59) |
| C5 | 32.3% (20 / 62) | 10.3% (7 / 68) | 2.9% (2 / 68) |
| TSLP | 96.3% (52 / 54) | 92.5% (62 / 67) | 0.0% (0 / 54) |
| IL17A | 7.9% (5 / 63) | 0.0% (0 / 65) | 0.0% (0 / 50) |
| FXI | 61.5% (24 / 39) | 20.9% (9 / 43) | 0.0% (0 / 33) |
| ACVR2B | 13.0% (10 / 77) | 5.9% (4 / 68) | 0.0% (0 / 66) |
| TNFRSF9 | 22.4% (15 / 67) | 24.4% (13 / 58) | 0.0% (0 / 59) |

**Table 4:** Fisher’s exact tests across antigens for IgDesign on HCDR3 and HCDR123 vs. SAbDab HCDR3 baseline. Significant *p*-values are bolded. At a significance level of *α* = 0.05 with *N* = 16 total tests, we require *p < α/N* = 0.05*/*16 = 0.003125 *≈* 3e-3 for a significant result. We note that IgDesign HCDR3 outperforms the baseline 8 out of 8 times and does so significantly 7 out of 8 times. IgDesign HCDR123 outperforms the baseline 7 out of 8 times and does so significantly 4 out of 8 times. We note that the baseline is only varying HCDR3 and keeping HCDR1 and HCDR2 fixed to the native sequence whereas IgDesign HCDR123 designs all three HCDRs.

| Antigen | IgDesign (HCDR3) |  | IgDesign (HCDR123) |  |
| --- | --- | --- | --- | --- |
| | Ratio to Baseline | $p$-value | Ratio to Baseline | $p$-value |
| CD40 | 19.6 | **5e-6** | 4.0 | 0.37 |
| IL36R | Inf | **5e-6** | Inf | **2.5e-4** |
| C5 | 11.0 | **5e-6** | 3.5 | 0.066 |
| TSLP | Inf | **0.0** | Inf | **0.0** |
| IL17A | Inf | 0.066 | N/A | 1.0 |
| FXI | Inf | **0.0** | Inf | 4.3e-3 |
| ACVR2B | Inf | **2e-3** | Inf | **2.6e-3** |
| TNFRSF9 | Inf | **4e-5** | Inf | **6e-5** |

**Figure 7:**
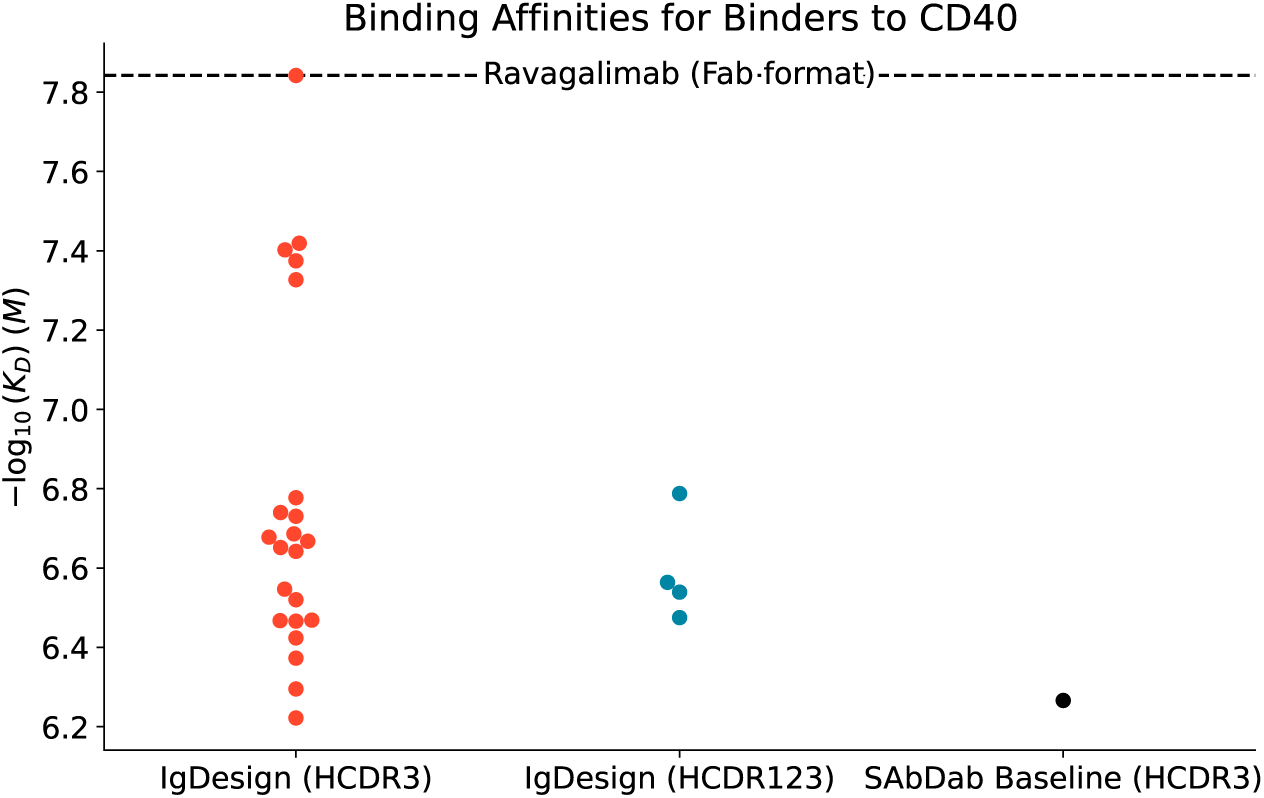
Binding affinities against CD40.

**Figure 8:**
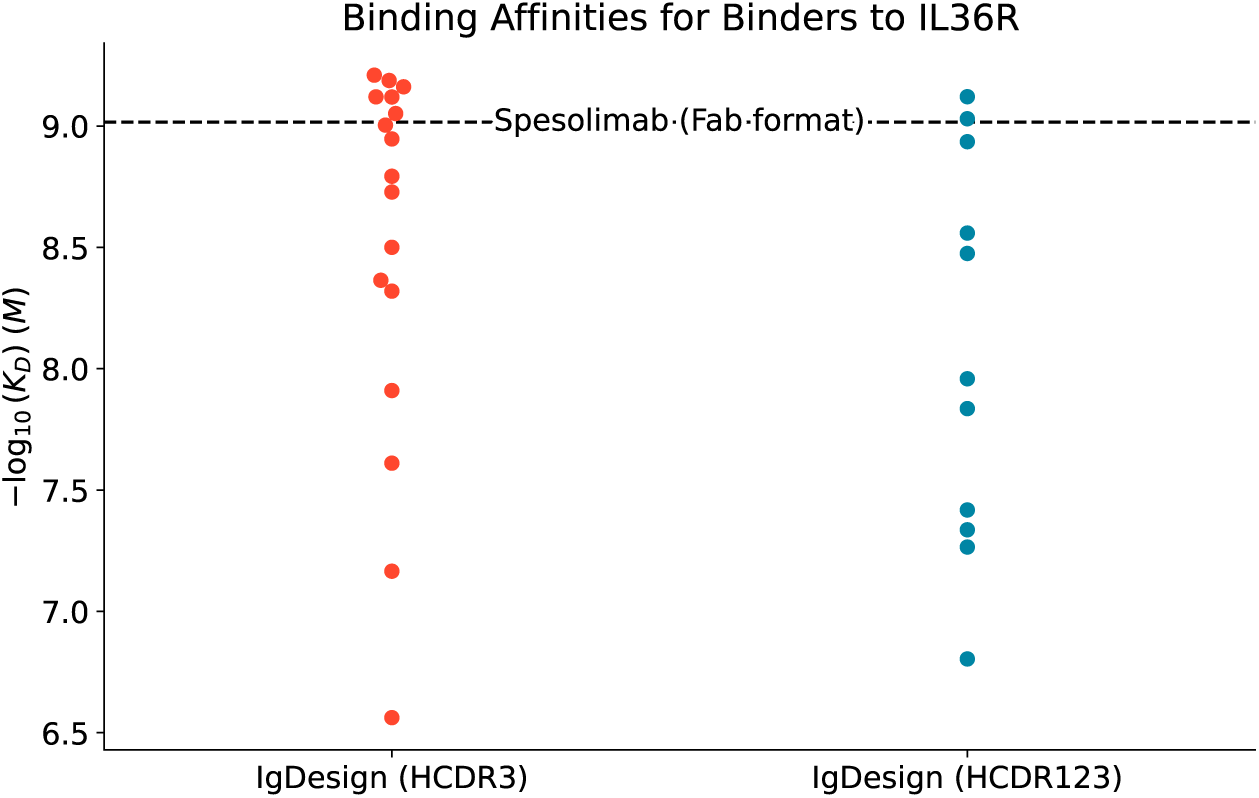
Binding affinities against IL36R.

**Figure 9:**
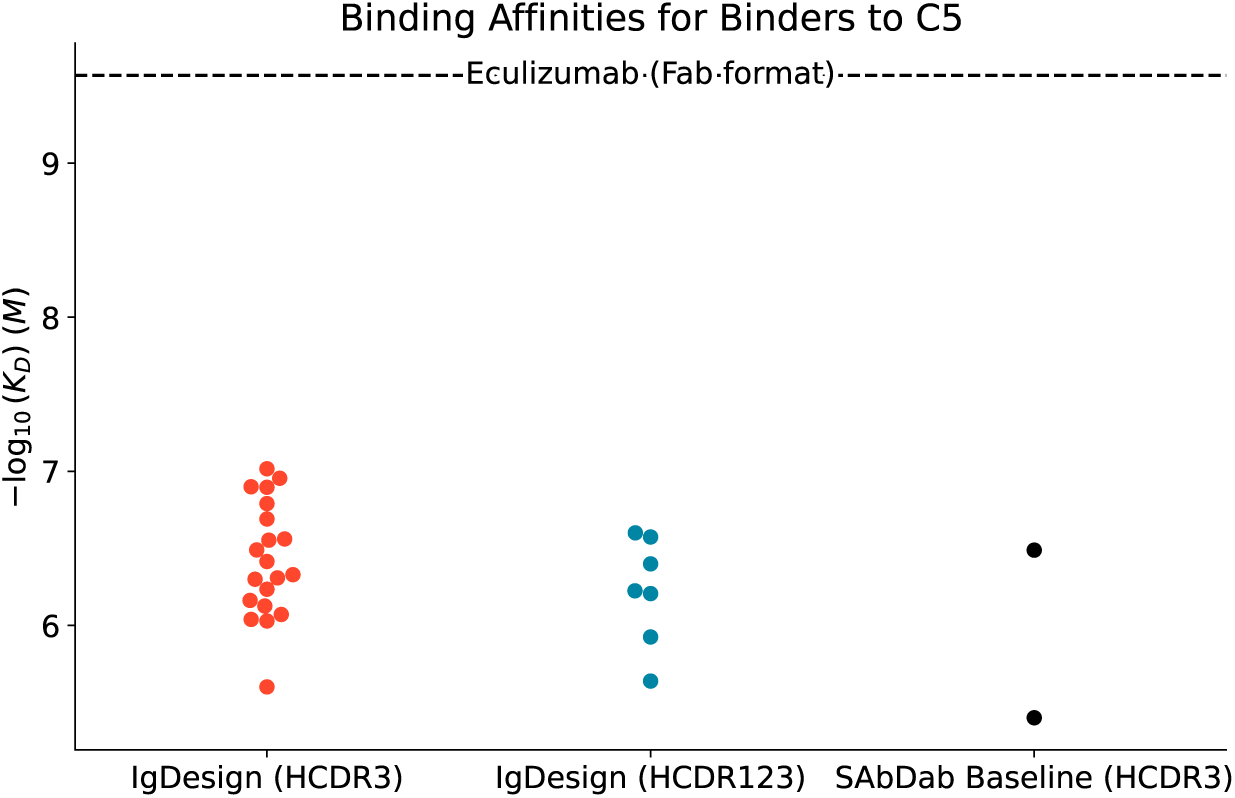
Binding affinities against C5.

**Figure 10:**
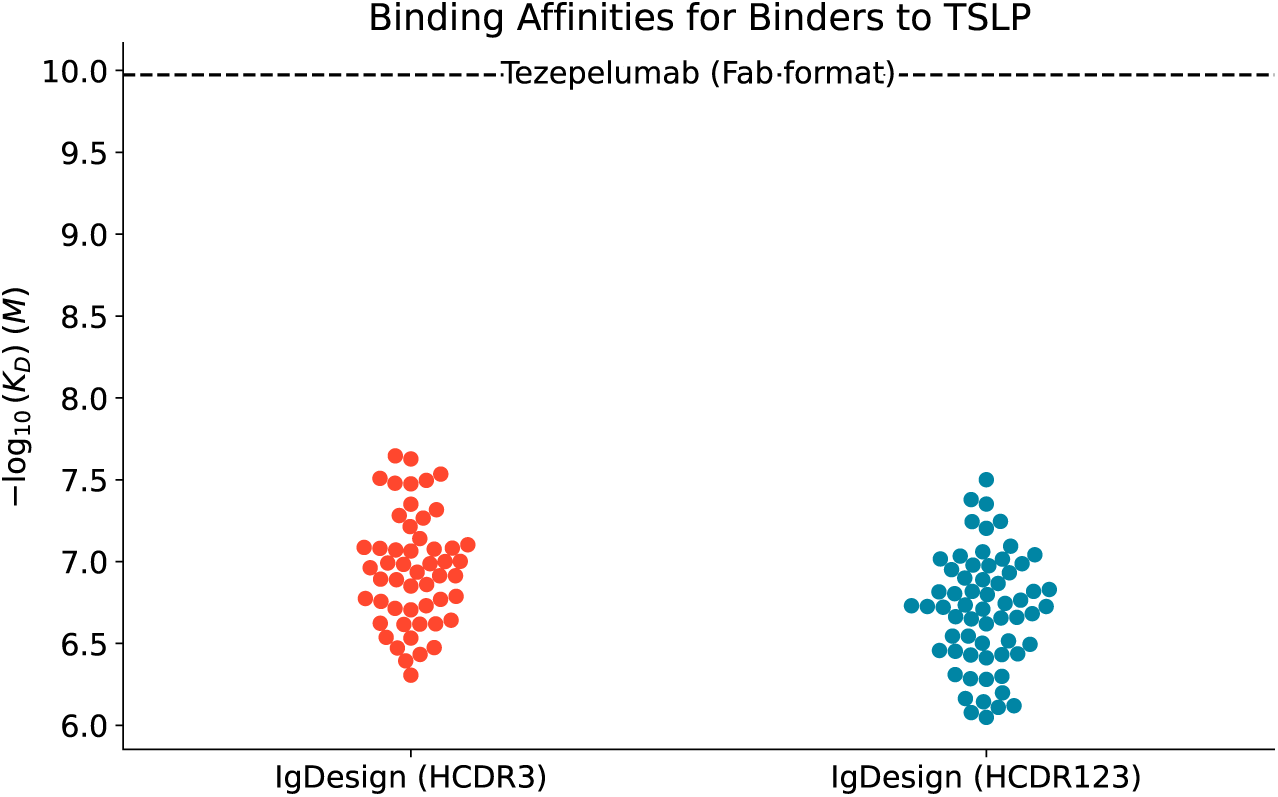
Binding affinities against TSLP.

**Figure 11:**
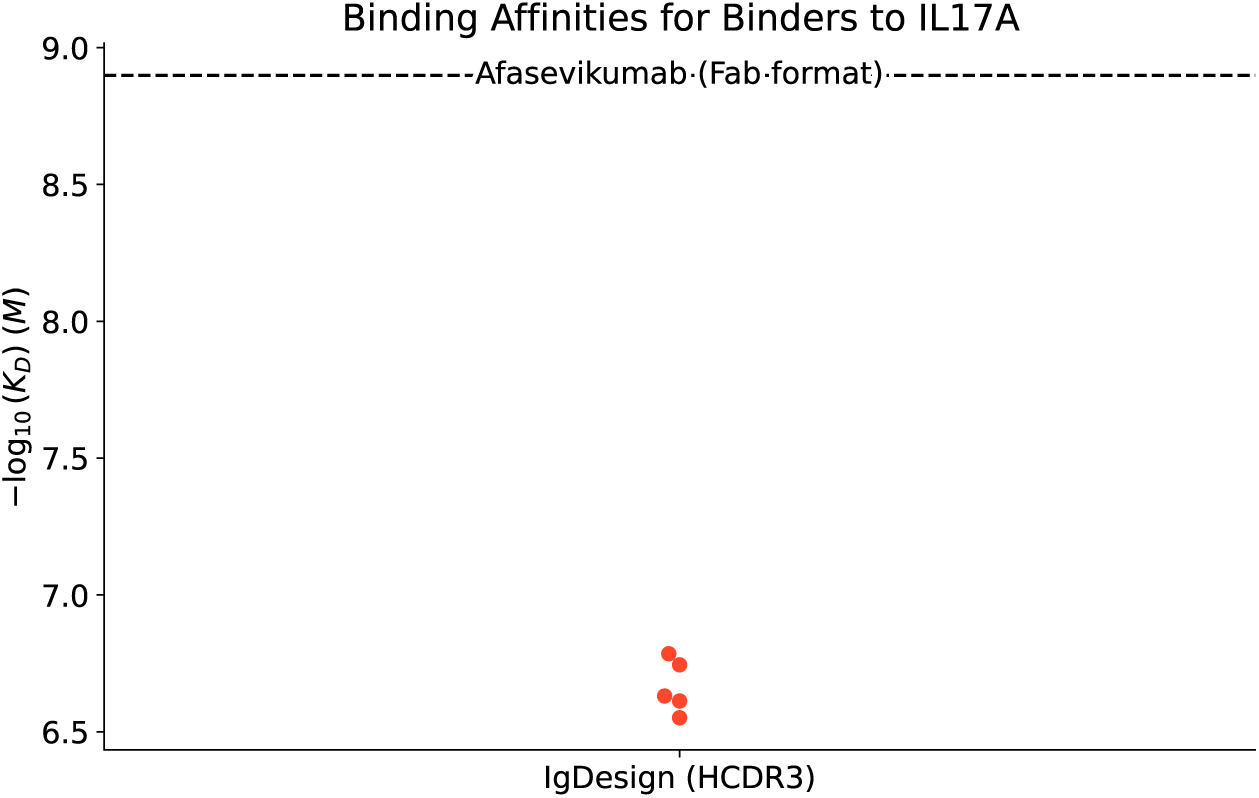
Binding affinities against IL17A.

**Figure 12:**
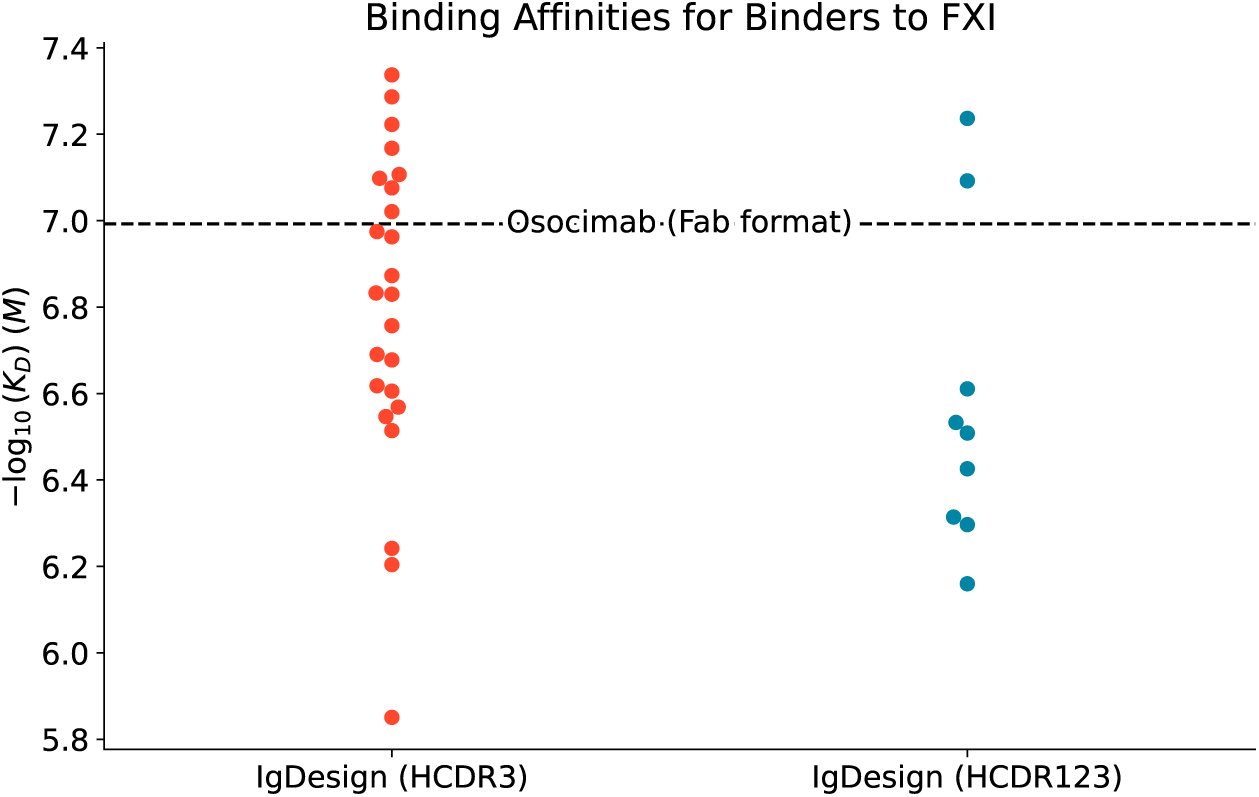
Binding affinities against FXI

**Figure 13:**
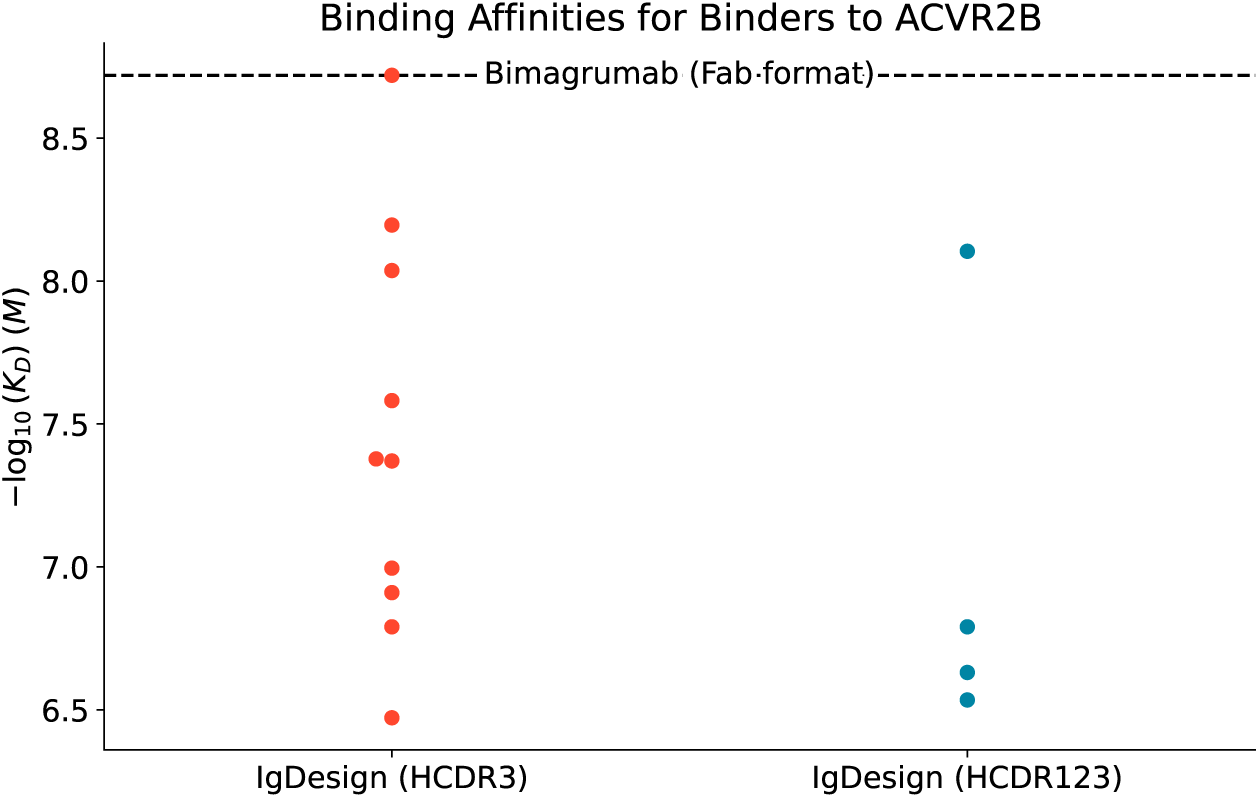
Binding affinities against ACVR2B.

**Figure 14:**
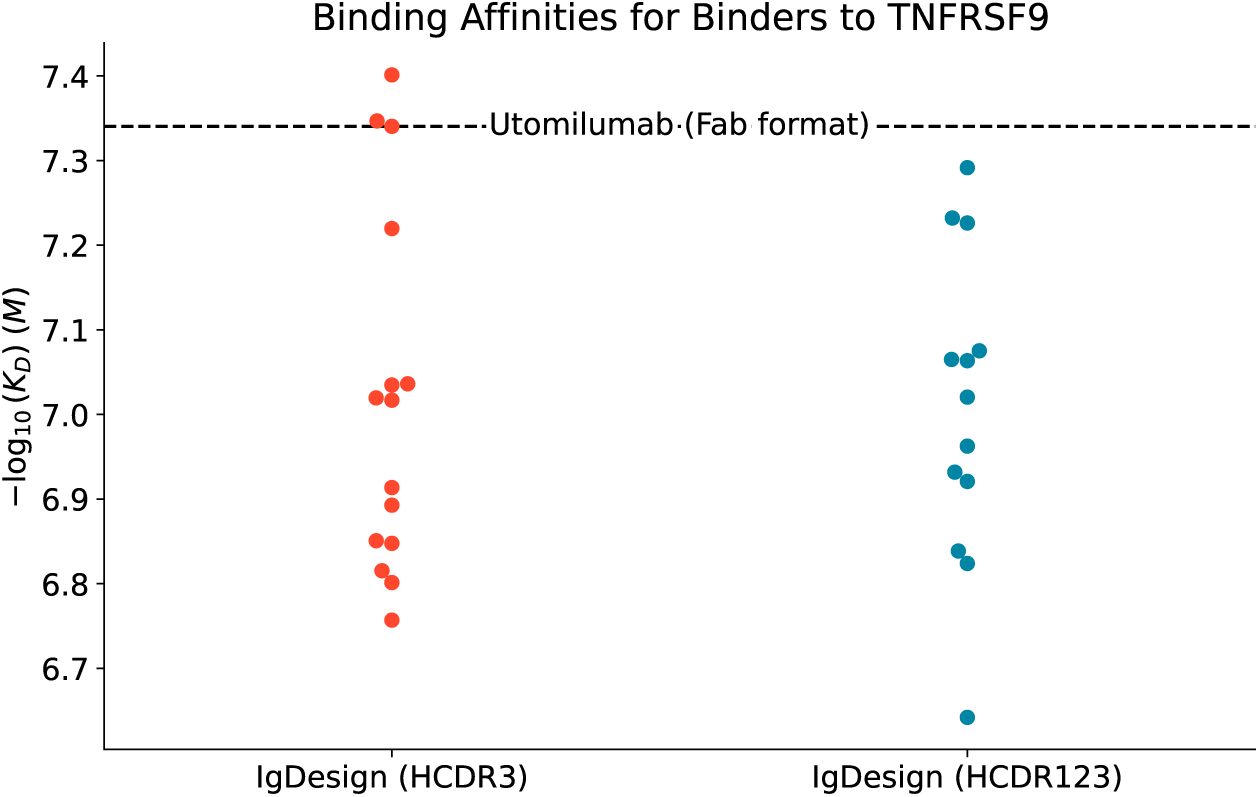
Binding affinities against TNFRSF9.

### Self-consistency RMSD (scRMSD)

We use the SPR datasets of binders and non-binders with affinities to assess the ability of the popular self-consistency RMSD (scRMSD) metric to discriminate between binders and non-binders and to predict affinity. We compute scRMSD across HCDR1, HCDR2, HCDR3, HCDR123, and the Fv (variable fragment) using ABodyBuilder2 [6], ABody-Builder3 [7], and ESMFold [8] as described in the Appendix. Full results, including visualizations of scRMSD distributions across antigens, design categories, and binding labels as well as tables of statistics can also be found in the Appendix.

In general, we see that scRMSD values are on average lower for binders than non-binders, with significance in some cases (Tables 6, 9, 12, 15, 18, 21, 24, 27), however we must consider possible sources of bias in the dataset. Firstly, there is risk of data leakage in the folding models and to assess this risk we compute scRMSD on the reference antibody, shown in Tables 5, 8, 11, 14, 17, 20, 23, 26. Data leakage could lead to bias towards the reference antibody sequence. Secondly, we see in Figures 15,16,17,18,19,20,21, 22 that the SAbDab HCDR3 scRMSD distribution is generally higher than that of inverse folding HCDR3s. While this does match with the measured binding rates, the scRMSD result is not surprising given that these baseline HCDR3s are sampled without conditioning on the HCDR3 loop structure and they may have been seen in a different antibody sequence during folding model training. When restricting to inverse folding designs only, the distribution of scRMSDs becomes tighter, in particular for HCDR3, and the metric has less discriminative power between binders and non-binders.

**Table 5:** scRMSD values of Ravagalimab (positive control). Predictions are generated using the unmodified antibody sequence and RMSDs are computed using the solved antibody structure.

| Model / Region | HCDR1 | HCDR2 | HCDR3 | HCDR123 | Fv |
| --- | --- | --- | --- | --- | --- |
| **ABodyBuilder2** | 0.41 | 0.60 | 0.36 | 0.46 | 0.37 |
| **ABodyBuilder3** | 0.45 | 0.93 | 0.62 | 0.69 | 0.56 |
| **ABodyBuilder3-LM** | 0.26 | 0.79 | 0.83 | 0.69 | 0.46 |
| **ESMFold** | 1.09 | 0.89 | 0.57 | 0.87 | 0.75 |

**Table 6:** Two-sided*t*-test values (*t, p*) between scRMSD values of Ravagalimab variants and binary binding labels to CD40. A negative *t*-statistic implies mean scRMSD values are lower for binders than non-binders.

| Model / Region | HCDR1 | HCDR2 | HCDR3 | HCDR123 | Fv |
| --- | --- | --- | --- | --- | --- |
| **ABodyBuilder2** | (-0.69, 0.49) | (-2.26, 0.025) | (-2.14, 0.034) | (-3.07, 0.0024) | (-3.49, 0.0006) |
| **ABodyBuilder3** | (-1.93, 0.056) | (-2.63, 0.0094) | (-1.51, 0.133) | (-3.39, 0.0009) | (-3.57, 0.0005) |
| **ABodyBuilder3-LM** | (-1.89, 0.061) | (-2.3, 0.0226) | (-2.03, 0.0439) | (-3.44, 0.0007) | (-3.26, 0.0013) |
| **ESMFold** | (-1.32, 0.19) | (-3.31, 0.0011) | (-3.29, 0.0235) | (-3.82, 0.0002) | (-2.6, 0.0102) |
| **Mean Ensemble** | (-2.31, 0.0218) | (-2.81, 0.0055) | (-2.14, 0.0337) | (-3.78, 0.0002) | (-3.69, 0.0003) |
| **Min Ensemble** | (-1.64, 0.1028) | (-2.5, 0.0134) | (-1.89, 0.0598) | (-3.55, 0.0005) | (-3.74, 0.0002) |
| **Max Ensemble** | (-1.34, 0.1804) | (-2.71, 0.0074) | (-1.98, 0.0492) | (-3.41, 0.0008) | (-2.7, 0.0075) |

**Table 7:** Pearson correlation (*n* = 27) between scRMSD values and affinities (*K_D_* (nM)) of Ravagalimab variant binders to CD40. A positive correlation implies lower scRMSD values correspond to lower *K_D_* values (higher affinities).

| Model / Region | HCDR1 | HCDR2 | HCDR3 | HCDR123 | Fv |
| --- | --- | --- | --- | --- | --- |
| **ABodyBuilder2** | -0.28 | 0.08 | 0.34 | 0.14 | 0.11 |
| **ABodyBuilder3** | -0.18 | 0.01 | 0.02 | 0.0 | -0.1 |
| **ABodyBuilder3-LM** | -0.33 | 0.02 | -0.22 | -0.03 | -0.05 |
| **ESMFold** | 0.29 | 0.02 | 0.22 | 0.09 | 0.07 |
| **Mean Ensemble** | -0.34 | 0.05 | 0.14 | 0.06 | 0.0 |
| **Min Ensemble** | -0.33 | 0.03 | 0.15 | 0.06 | 0.08 |
| **Max Ensemble** | 0.29 | 0.06 | -0.02 | 0.04 | 0.07 |

**Table 8:** scRMSD values of Spesolimab (positive control). Predictions are generated using the unmodified antibody sequence and RMSDs are computed using the solved antibody structure.

| Model / Region | HCDR1 | HCDR2 | HCDR3 | HCDR123 | Fv |
| --- | --- | --- | --- | --- | --- |
| **ABodyBuilder2** | 0.62 | 0.3 | 0.66 | 0.57 | 0.38 |
| **ABodyBuilder3** | 0.63 | 0.4 | 1.14 | 0.84 | 0.55 |
| **ABodyBuilder3-LM** | 1.19 | 0.71 | 1.21 | 1.09 | 0.65 |
| **ESMFold** | 1.02 | 0.65 | 3.35 | 2.28 | 1.18 |

**Table 9:** *t*-test values (*t, p*) between scRMSD values of Spesolimab variants and binary binding labels to IL36R. A negative *t*-statistic implies mean scRMSD values are lower for binders than non-binders.

| Model / Region | HCDR1 | HCDR2 | HCDR3 | HCDR123 | Fv |
| --- | --- | --- | --- | --- | --- |
| **ABodyBuilder2** | (0.6, 0.551) | (-1.21, 0.2271) | (-2.05, 0.0421) | (-1.91, 0.0576) | (-1.95, 0.0526) |
| **ABodyBuilder3** | (-0.64, 0.5212) | (0.42, 0.6729) | (-3.03, 0.0028) | (-2.83, 0.0052) | (-2.28, 0.0237) |
| **ABodyBuilder3-LM** | (0.24, 0.8102) | (-0.49, 0.6227) | (-2.24, 0.0266) | (-2.03, 0.0437) | (-1.08, 0.2811) |
| **ESMFold** | (0.59, 0.5589) | (-0.88, 0.3805) | (-2.84, 0.005) | (-2.77, 0.0063) | (-2.45, 0.0152) |
| **Mean Ensemble** | (0.19, 0.852) | (-0.76, 0.4483) | (-3.48, 0.0006) | (-3.27, 0.0013) | (-2.75, 0.0066) |
| **Min Ensemble** | (2.0, 0.0475) | (-1.42, 0.1585) | (-3.46, 0.0007) | (-3.26, 0.0013) | (-1.88, 0.0613) |
| **Max Ensemble** | (0.26, 0.7966) | (-0.72, 0.4712) | (-2.88, 0.0044) | (-2.58, 0.0108) | (-2.54, 0.012) |

**Table 10:**
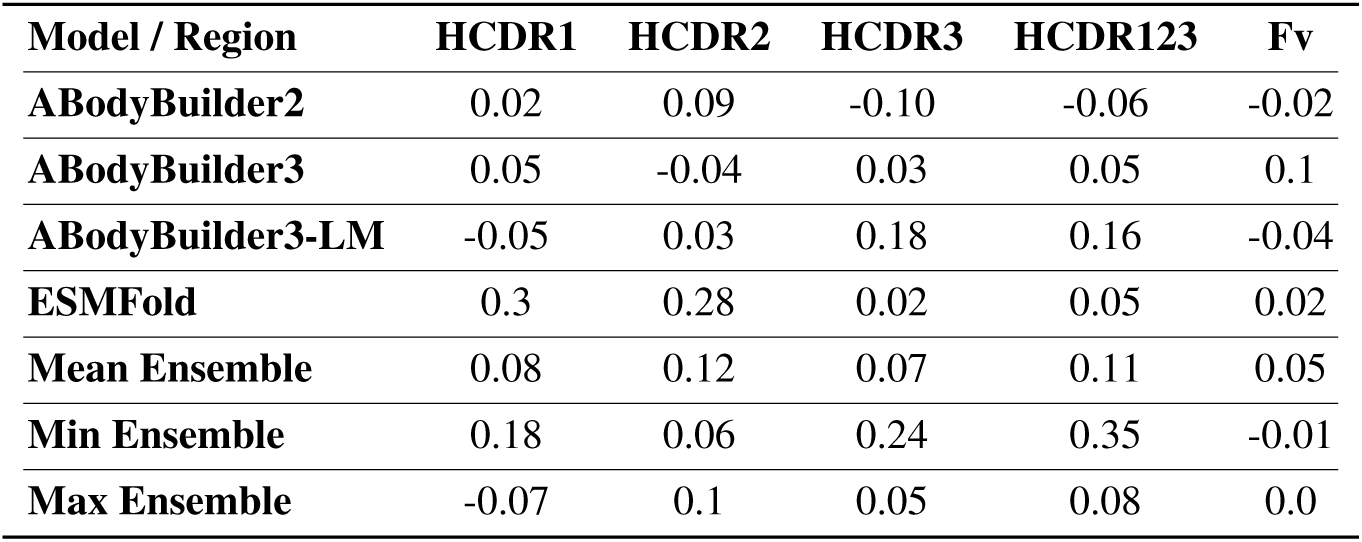
Pearson correlation (*n* = 29) between scRMSD values and affinities (*K_D_* (nM)) of Spesolimab variant binders to IL36R. A positive correlation implies lower scRMSD values correspond to lower *K_D_* values (higher affinities).

| Model / Region | HCDR1 | HCDR2 | HCDR3 | HCDR123 | Fv |
| --- | --- | --- | --- | --- | --- |
| **ABodyBuilder2** | 0.02 | 0.09 | -0.10 | -0.06 | -0.02 |
| **ABodyBuilder3** | 0.05 | -0.04 | 0.03 | 0.05 | 0.1 |
| **ABodyBuilder3-LM** | -0.05 | 0.03 | 0.18 | 0.16 | -0.04 |
| **ESMFold** | 0.3 | 0.28 | 0.02 | 0.05 | 0.02 |
| **Mean Ensemble** | 0.08 | 0.12 | 0.07 | 0.11 | 0.05 |
| **Min Ensemble** | 0.18 | 0.06 | 0.24 | 0.35 | -0.01 |
| **Max Ensemble** | -0.07 | 0.1 | 0.05 | 0.08 | 0.0 |

**Table 11:** scRMSD values of Eculizumab (positive control). Predictions are generated using the unmodified antibody sequence and RMSDs are computed using the solved antibody structure.

| Model / Region | HCDR1 | HCDR2 | HCDR3 | HCDR123 | Fv |
| --- | --- | --- | --- | --- | --- |
| **ABodyBuilder2** | 1.15 | 1.08 | 2.23 | 1.75 | 1.3 |
| **ABodyBuilder3** | 1.32 | 1.14 | 2.4 | 1.89 | 1.3 |
| **ABodyBuilder3-LM** | 1.2 | 1.2 | 1.84 | 1.54 | 1.24 |
| **ESMFold** | 0.85 | 1.07 | 1.66 | 1.35 | 1.2 |

**Table 12:** *t*-test values (*t, p*) between scRMSD values of Eculizumab variants and binary binding labels to C5. A negative *t*-statistic implies mean scRMSD values are lower for binders than non-binders.

| Model / Region | HCDR1 | HCDR2 | HCDR3 | HCDR123 | Fv |
| --- | --- | --- | --- | --- | --- |
| **ABodyBuilder2** | (1.69, 0.0927) | (0.64, 0.525) | (-1.94, 0.054) | (-1.71, 0.0883) | (-0.65, 0.5168) |
| **ABodyBuilder3** | (0.33, 0.7412) | (-0.2, 0.8451) | (0.34, 0.7342) | (0.31, 0.7532) | (-0.17, 0.8629) |
| **ABodyBuilder3-LM** | (-2.34, 0.0201) | (-1.8, 0.074) | (-2.53, 0.0122) | (-2.6, 0.01) | (-2.75, 0.0065) |
| **ESMFold** | (-3.65, 0.0003) | (-0.94, 0.3481) | (-4.5, 0.0) | (-4.13, 0.0001) | (-3.88, 0.0001) |
| **Mean Ensemble** | (-0.58, 0.5621) | (-1.14, 0.2562) | (-3.1, 0.0022) | (-2.97, 0.0033) | (-2.65, 0.0088) |
| **Min Ensemble** | (-2.95, 0.0035) | (-1.42, 0.1572) | (-5.29, 0.0) | (-4.64, 0.0) | (-4.36, 0.0) |
| **Max Ensemble** | (0.24, 0.807) | (-1.23, 0.2195) | (-1.79, 0.075) | (-1.87, 0.0625) | (-2.19, 0.0301) |

**Table 13:** Pearson correlation (*n* = 30) between scRMSD values and affinities (*K_D_* (nM)) of Eculizumab variant binders to C5. A positive correlation implies lower scRMSD values correspond to lower *K_D_* values (higher affinities).

| Model / Region | HCDR1 | HCDR2 | HCDR3 | HCDR123 | Fv |
| --- | --- | --- | --- | --- | --- |
| **ABodyBuilder2** | 0.15 | 0.17 | 0.4 | 0.39 | 0.32 |
| **ABodyBuilder3** | -0.04 | 0.24 | -0.05 | -0.04 | 0.05 |
| **ABodyBuilder3-LM** | 0.43 | 0.31 | 0.46 | 0.48 | 0.53 |
| **ESMFold** | 0.41 | 0.1 | 0.58 | 0.58 | 0.62 |
| **Mean Ensemble** | 0.3 | 0.38 | 0.38 | 0.39 | 0.41 |
| **Min Ensemble** | 0.41 | 0.37 | 0.61 | 0.52 | 0.64 |
| **Max Ensemble** | 0.07 | 0.28 | 0.26 | 0.26 | 0.27 |

**Table 14:** scRMSD values of Tezepelumab (positive control). Predictions are generated using the unmodified antibody sequence and RMSDs are computed using the solved antibody structure.

| Model / Region | HCDR1 | HCDR2 | HCDR3 | HCDR123 | Fv |
| --- | --- | --- | --- | --- | --- |
| **ABodyBuilder2** | 0.31 | 0.52 | 0.58 | 0.51 | 0.37 |
| **ABodyBuilder3** | 0.63 | 0.65 | 2.22 | 1.61 | 0.75 |
| **ABodyBuilder3-LM** | 0.84 | 0.67 | 2.33 | 1.71 | 0.76 |
| **ESMFold** | 0.71 | 1.13 | 4.68 | 3.32 | 1.36 |

**Table 15:**
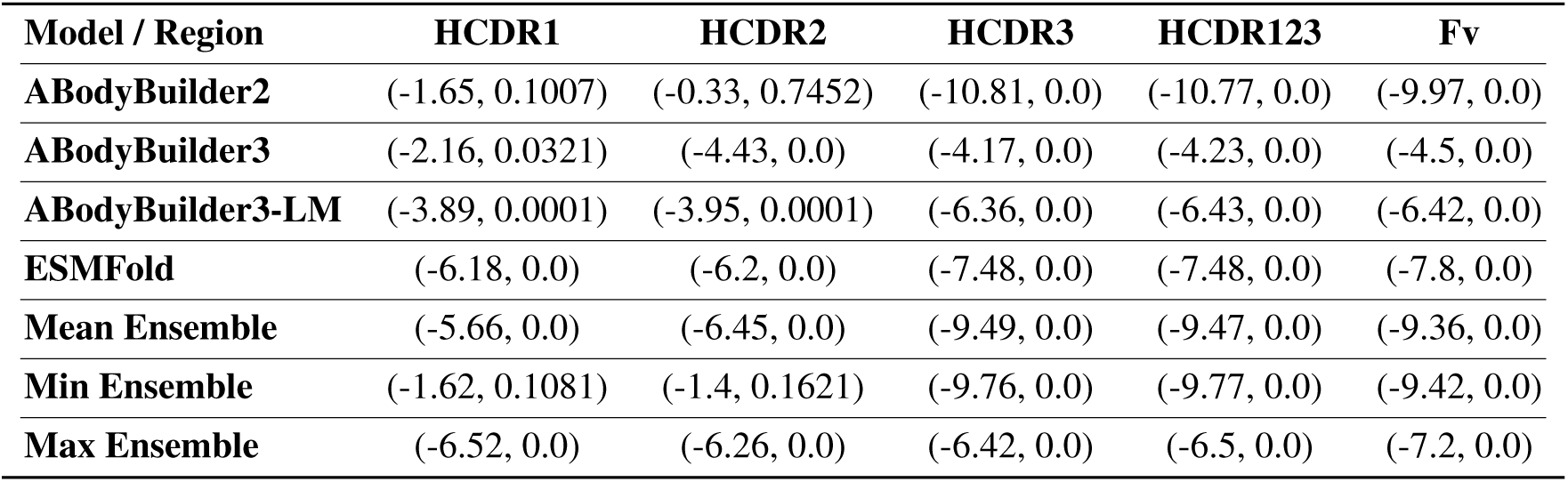
*t*-test values (*t, p*) between scRMSD values of Tezepelumab variants and binary binding labels to TSLP. A negative *t*-statistic implies mean scRMSD values are lower for binders than non-binders.

| Model / Region | HCDR1 | HCDR2 | HCDR3 | HCDR123 | Fv |
| --- | --- | --- | --- | --- | --- |
| **ABodyBuilder2** | (-1.65, 0.1007) | (-0.33, 0.7452) | (-10.81, 0.0) | (-10.77, 0.0) | (-9.97, 0.0) |
| **ABodyBuilder3** | (-2.16, 0.0321) | (-4.43, 0.0) | (-4.17, 0.0) | (-4.23, 0.0) | (-4.5, 0.0) |
| **ABodyBuilder3-LM** | (-3.89, 0.0001) | (-3.95, 0.0001) | (-6.36, 0.0) | (-6.43, 0.0) | (-6.42, 0.0) |
| **ESMFold** | (-6.18, 0.0) | (-6.2, 0.0) | (-7.48, 0.0) | (-7.48, 0.0) | (-7.8, 0.0) |
| **Mean Ensemble** | (-5.66, 0.0) | (-6.45, 0.0) | (-9.49, 0.0) | (-9.47, 0.0) | (-9.36, 0.0) |
| **Min Ensemble** | (-1.62, 0.1081) | (-1.4, 0.1621) | (-9.76, 0.0) | (-9.77, 0.0) | (-9.42, 0.0) |
| **Max Ensemble** | (-6.52, 0.0) | (-6.26, 0.0) | (-6.42, 0.0) | (-6.5, 0.0) | (-7.2, 0.0) |

**Table 16:** Pearson correlation (*n* = 115) between scRMSD values and affinities (*K_D_* (nM)) of Tezepelumab variant binders to TSLP. A positive correlation implies lower scRMSD values correspond to lower *K_D_* values (higher affinities).

| Model / Region | HCDR1 | HCDR2 | HCDR3 | HCDR123 | Fv |
| --- | --- | --- | --- | --- | --- |
| **ABodyBuilder2** | 0.26 | 0.16 | 0.09 | 0.11 | 0.22 |
| **ABodyBuilder3** | -0.02 | -0.13 | -0.11 | -0.11 | -0.12 |
| **ABodyBuilder3-LM** | -0.04 | -0.11 | -0.15 | -0.15 | -0.14 |
| **ESMFold** | 0.01 | -0.24 | 0.09 | 0.09 | 0.08 |
| **Mean Ensemble** | 0.1 | -0.11 | -0.02 | -0.02 | 0.02 |
| **Min Ensemble** | 0.15 | -0.06 | 0.12 | 0.14 | 0.2 |
| **Max Ensemble** | -0.01 | -0.16 | -0.07 | -0.07 | -0.04 |

**Table 17:** scRMSD values of Afasevikumab (positive control). Predictions are generated using the unmodified antibody sequence and RMSDs are computed using the solved antibody structure.

| Model / Region | HCDR1 | HCDR2 | HCDR3 | HCDR123 | Fv |
| --- | --- | --- | --- | --- | --- |
| **ABodyBuilder2** | 0.41 | 0.45 | 0.5 | 0.47 | 0.41 |
| **ABodyBuilder3** | 0.46 | 0.4 | 1.57 | 1.15 | 0.71 |
| **ABodyBuilder3-LM** | 0.54 | 0.47 | 1.13 | 0.88 | 0.68 |
| **ESMFold** | 1.02 | 2.61 | 3.62 | 2.92 | 1.56 |

**Table 18:** *t*-test values (*t, p*) between scRMSD values of Afasevikumab variants and binary binding labels to IL17A. A negative *t*-statistic implies mean scRMSD values are lower for binders than non-binders.

| Model / Region | HCDR1 | HCDR2 | HCDR3 | HCDR123 | Fv |
| --- | --- | --- | --- | --- | --- |
| **ABodyBuilder2** | (-0.13, 0.8935) | (0.1, 0.9166) | (-2.57, 0.0109) | (-2.52, 0.0126) | (-2.59, 0.0103) |
| **ABodyBuilder3** | (-0.06, 0.9552) | (-2.31, 0.0219) | (-1.29, 0.1985) | (-1.34, 0.1827) | (-1.21, 0.2296) |
| **ABodyBuilder3-LM** | (-1.09, 0.2773) | (-2.05, 0.0418) | (-1.76, 0.0801) | (-1.8, 0.0738) | (-1.8, 0.0742) |
| **ESMFold** | (-0.53, 0.5951) | (1.14, 0.256) | (-0.29, 0.7723) | (-0.11, 0.9091) | (-0.18, 0.8596) |
| **Mean Ensemble** | (-0.69, 0.4884) | (0.3, 0.7645) | (-1.71, 0.0888) | (-1.65, 0.1008) | (-1.65, 0.1016) |
| **Min Ensemble** | (-0.83, 0.4064) | (-1.85, 0.0659) | (-2.18, 0.0305) | (-2.23, 0.0272) | (-2.71, 0.0073) |
| **Max Ensemble** | (-0.54, 0.5919) | (1.13, 0.2608) | (-0.57, 0.567) | (-0.3, 0.7664) | (-0.18, 0.8596) |

**Figure 15:**
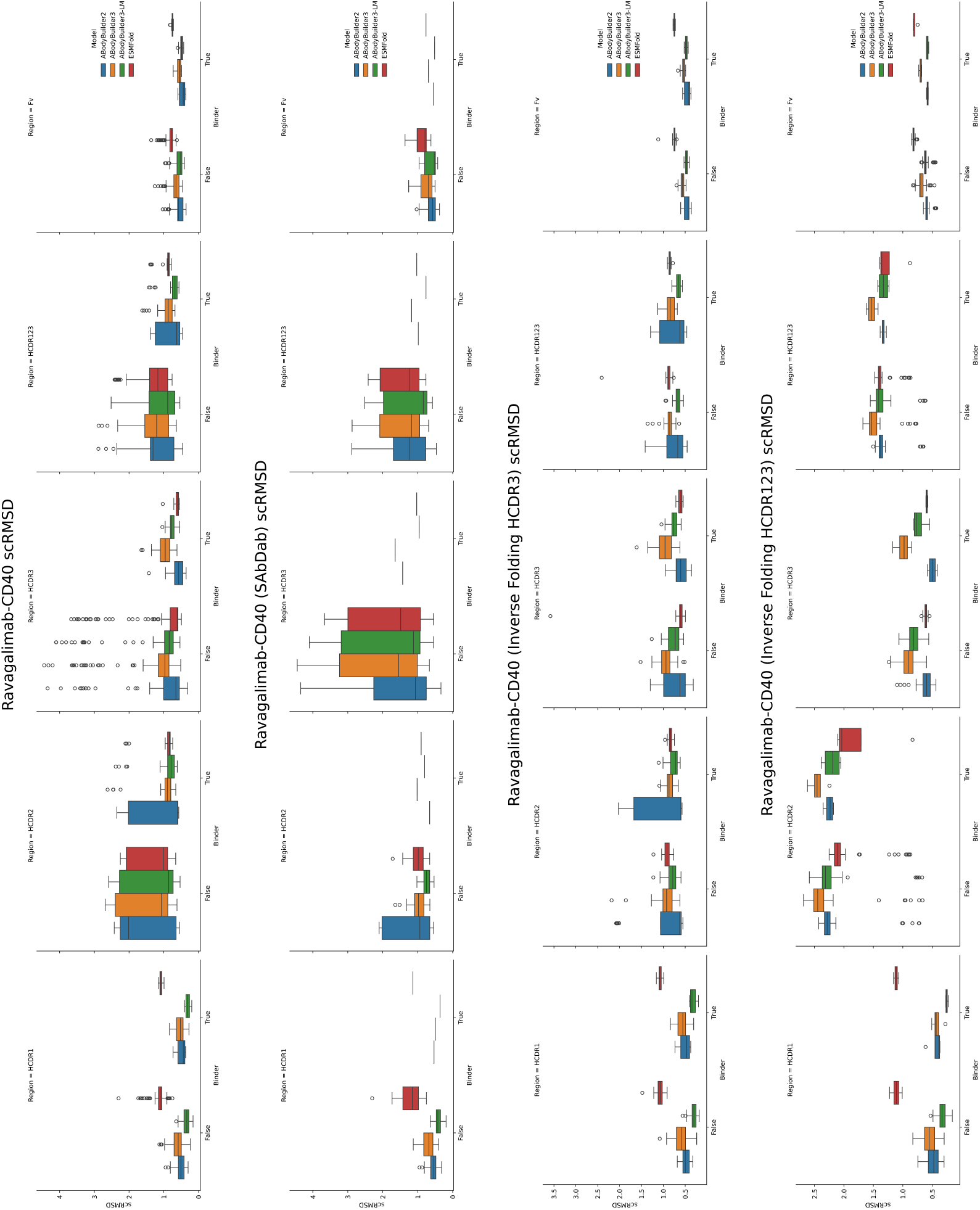
scRMSD distributions of Ravagalimab variants. Values for each of 5 regions (HCDR1, HCDR2, HCDR3, HCDR123, and Fv) and 4 models (ABodyBuilder2, ABodyBuilder3, ABodyBuilder3-LM, and ESMFold) are shown. Distributions are also broken down by binary binding labels and shown (from top to bottom) for all sequences, sequences from the SAbDab baseline, inverse folding HCDR3 designs, and inverse folding HCDR123 designs.

**Figure 16:**
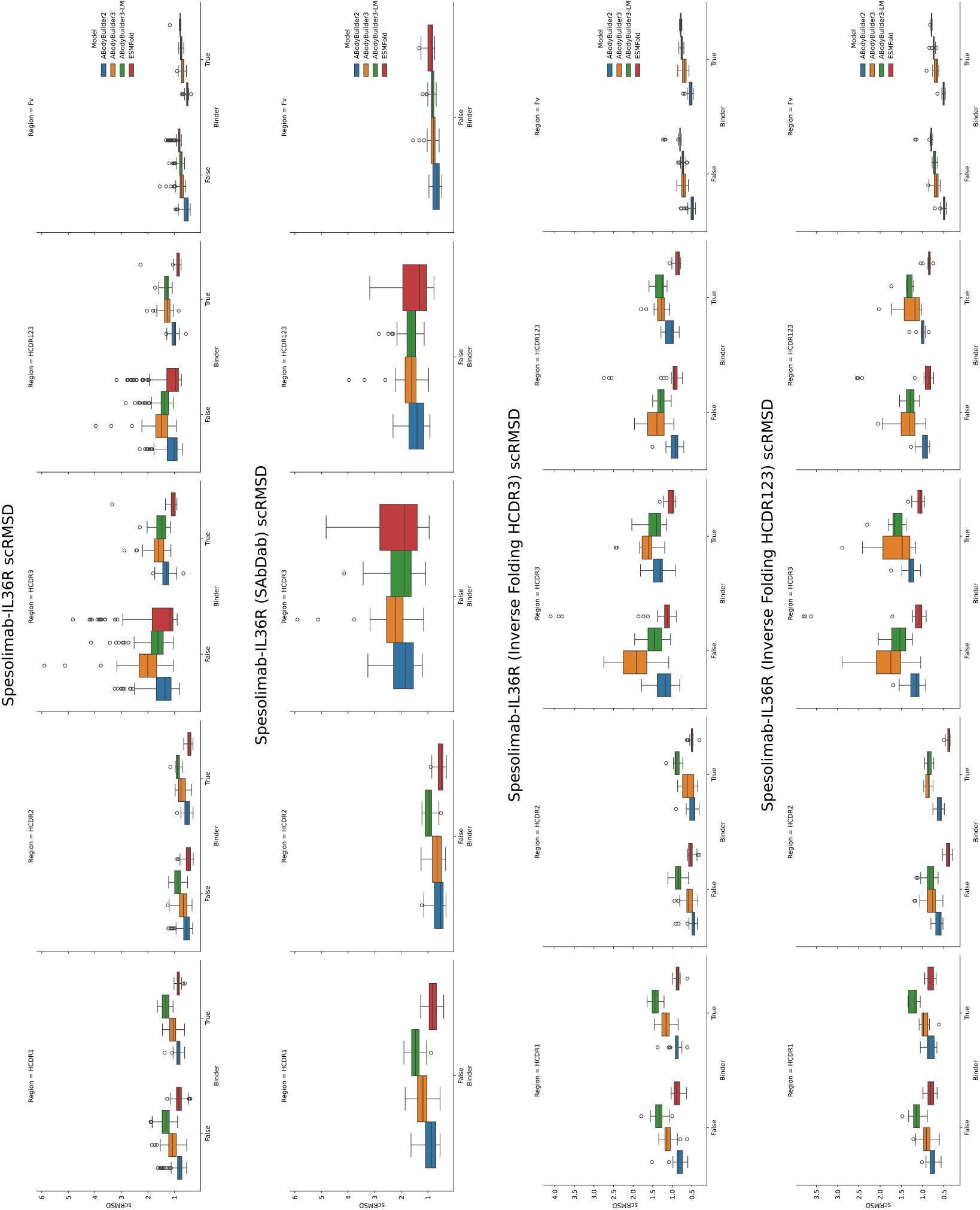
scRMSD distributions of Spesolimab variants. Values for each of 5 regions (HCDR1, HCDR2, HCDR3, HCDR123, and Fv) and 4 models (ABodyBuilder2, ABodyBuilder3, ABodyBuilder3-LM, and ESMFold) are shown. Distributions are also broken down by binary binding labels and shown (from top to bottom) for all sequences, sequences from the SAbDab baseline, inverse folding HCDR3 designs, and inverse folding HCDR123 designs.

**Figure 17:**
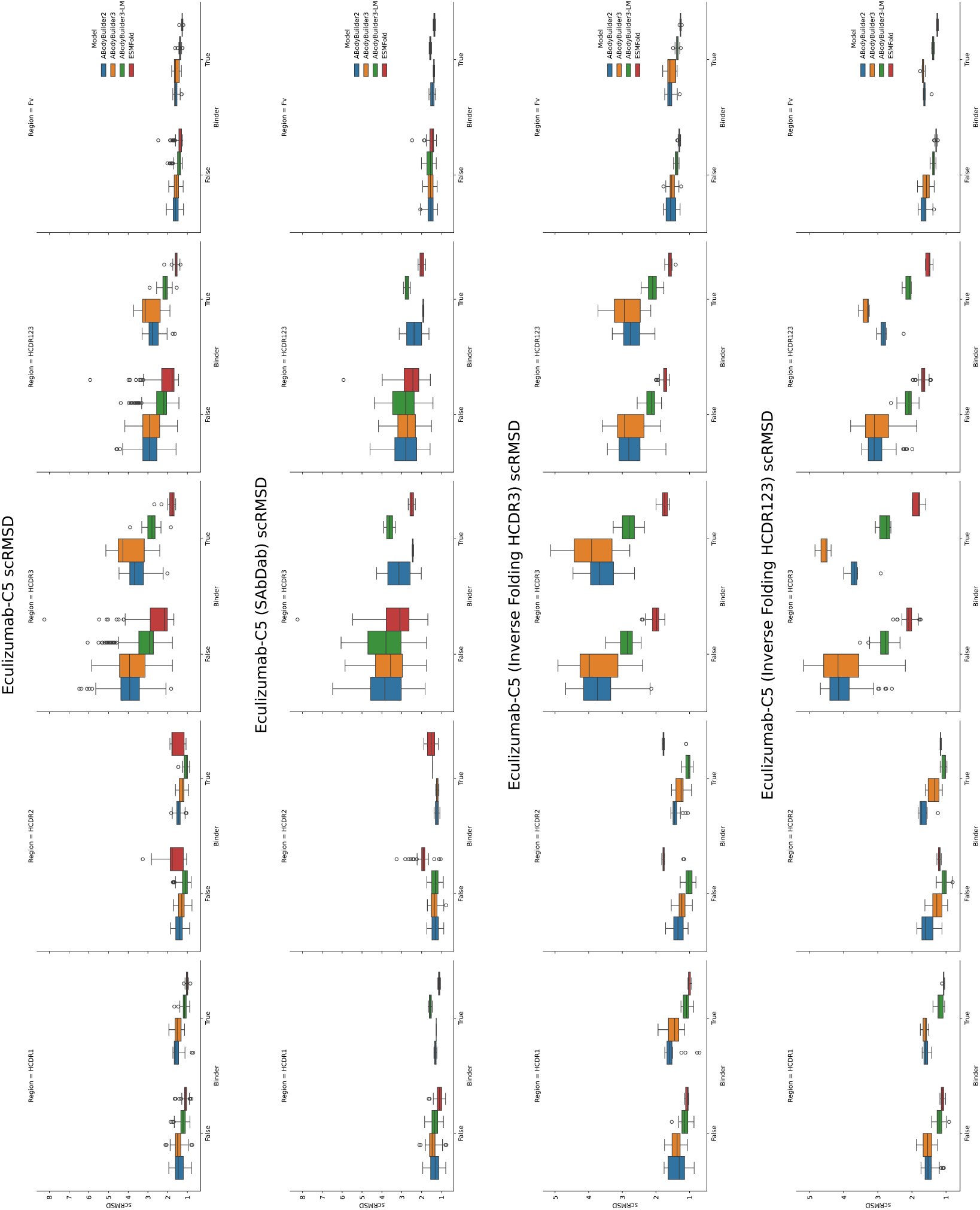
scRMSD distributions of Eculizumab variants. Values for each of 5 regions (HCDR1, HCDR2, HCDR3, HCDR123, and Fv) and 4 models (ABodyBuilder2, ABodyBuilder3, ABodyBuilder3-LM, and ESMFold) are shown. Distributions are also broken down by binary binding labels and shown (from top to bottom) for all sequences, sequences from the SAbDab baseline, inverse folding HCDR3 designs, and inverse folding HCDR123 designs.

**Figure 18:**
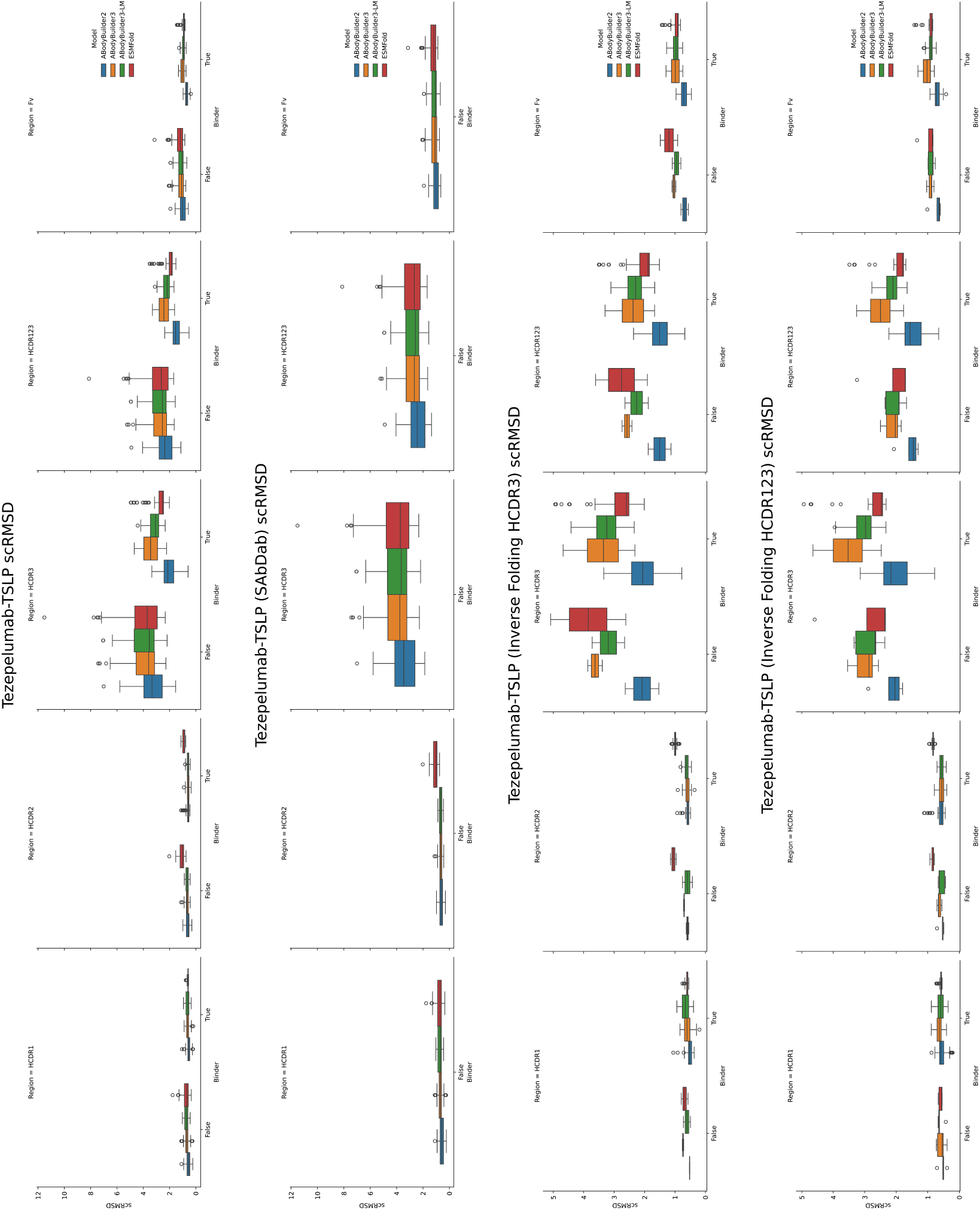
scRMSD distributions of Tezepelumab variants. Values for each of 5 regions (HCDR1, HCDR2, HCDR3, HCDR123, and Fv) and 4 models (ABodyBuilder2, ABodyBuilder3, ABodyBuilder3-LM, and ESMFold) are shown. Distributions are also broken down by binary binding labels and shown (from top to bottom) for all sequences, sequences from the SAbDab baseline, inverse folding HCDR3 designs, and inverse folding HCDR123 designs.

**Figure 19:**
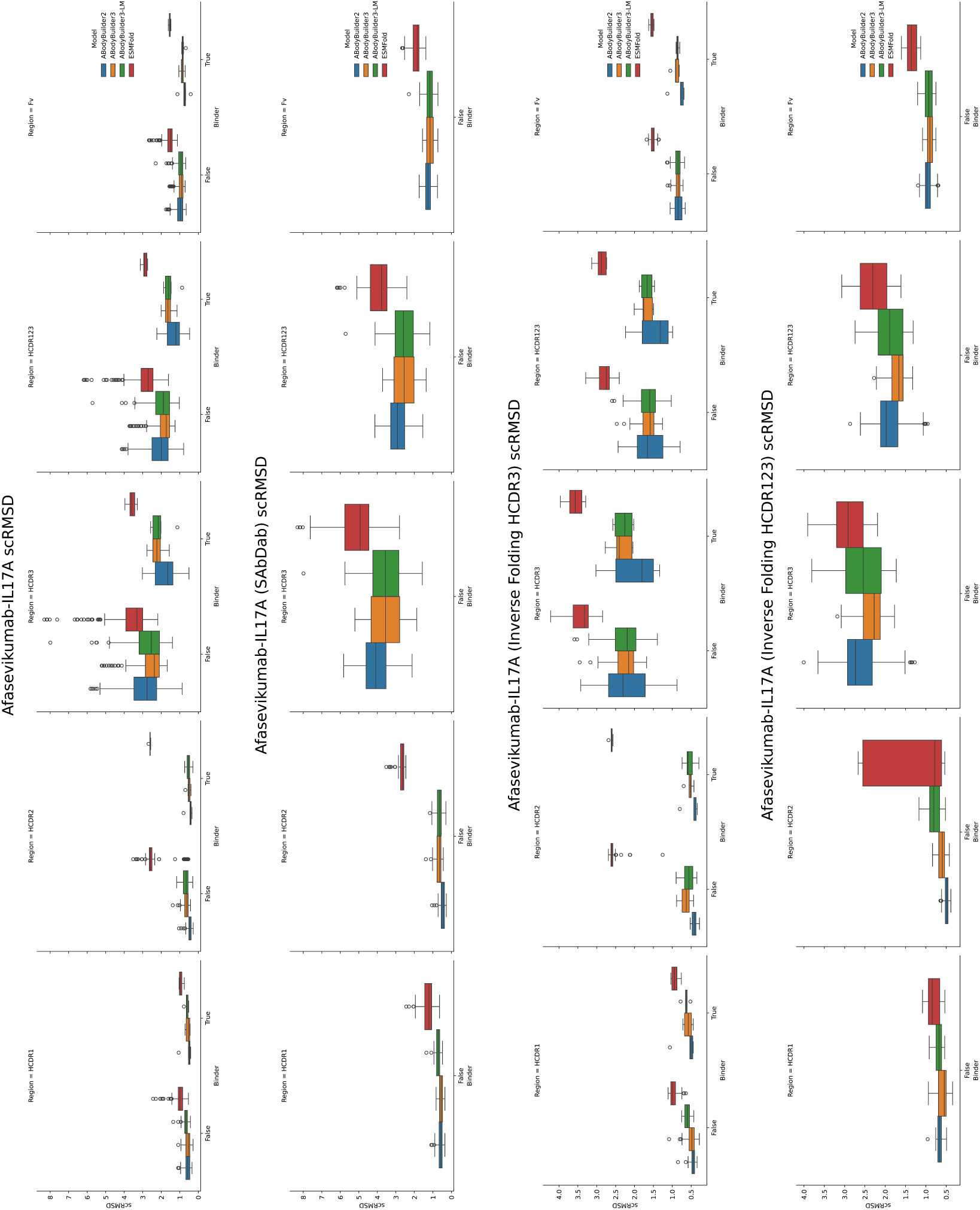
scRMSD distributions of Afasevikumab variants. Values for each of 5 regions (HCDR1, HCDR2, HCDR3, HCDR123, and Fv) and 4 models (ABodyBuilder2, ABodyBuilder3, ABodyBuilder3-LM, and ESMFold) are shown. Distributions are also broken down by binary binding labels and shown (from top to bottom) for all sequences, sequences from the SAbDab baseline, inverse folding HCDR3 designs, and inverse folding HCDR123 designs.

**Figure 20:**
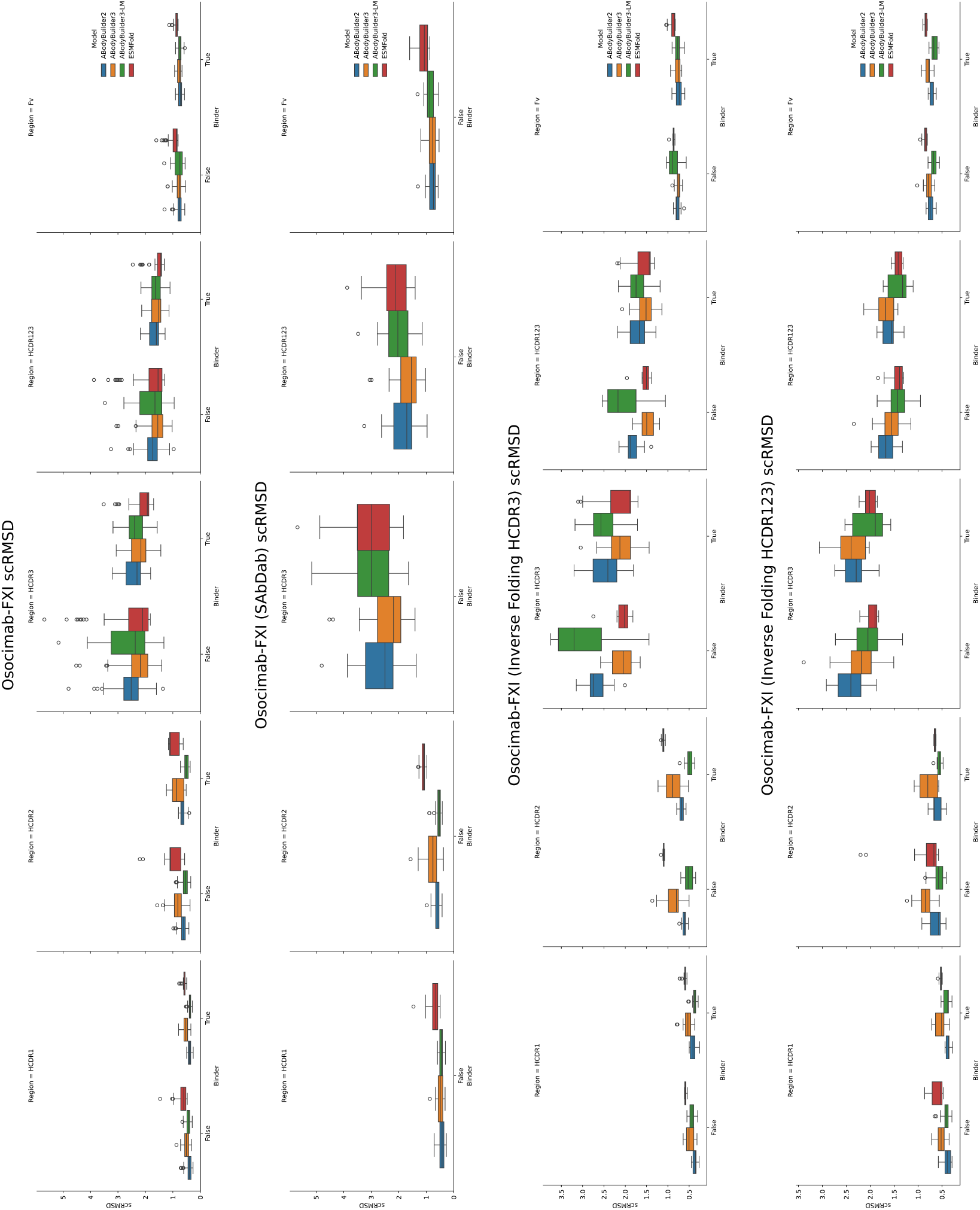
scRMSD distributions of Osocimab variants. Values for each of 5 regions (HCDR1, HCDR2, HCDR3, HCDR123, and Fv) and 4 models (ABodyBuilder2, ABodyBuilder3, ABodyBuilder3-LM, and ESMFold) are shown. Distributions are also broken down by binary binding labels and shown (from top to bottom) for all sequences, sequences from the SAbDab baseline, inverse folding HCDR3 designs, and inverse folding HCDR123 designs.

**Figure 21:**
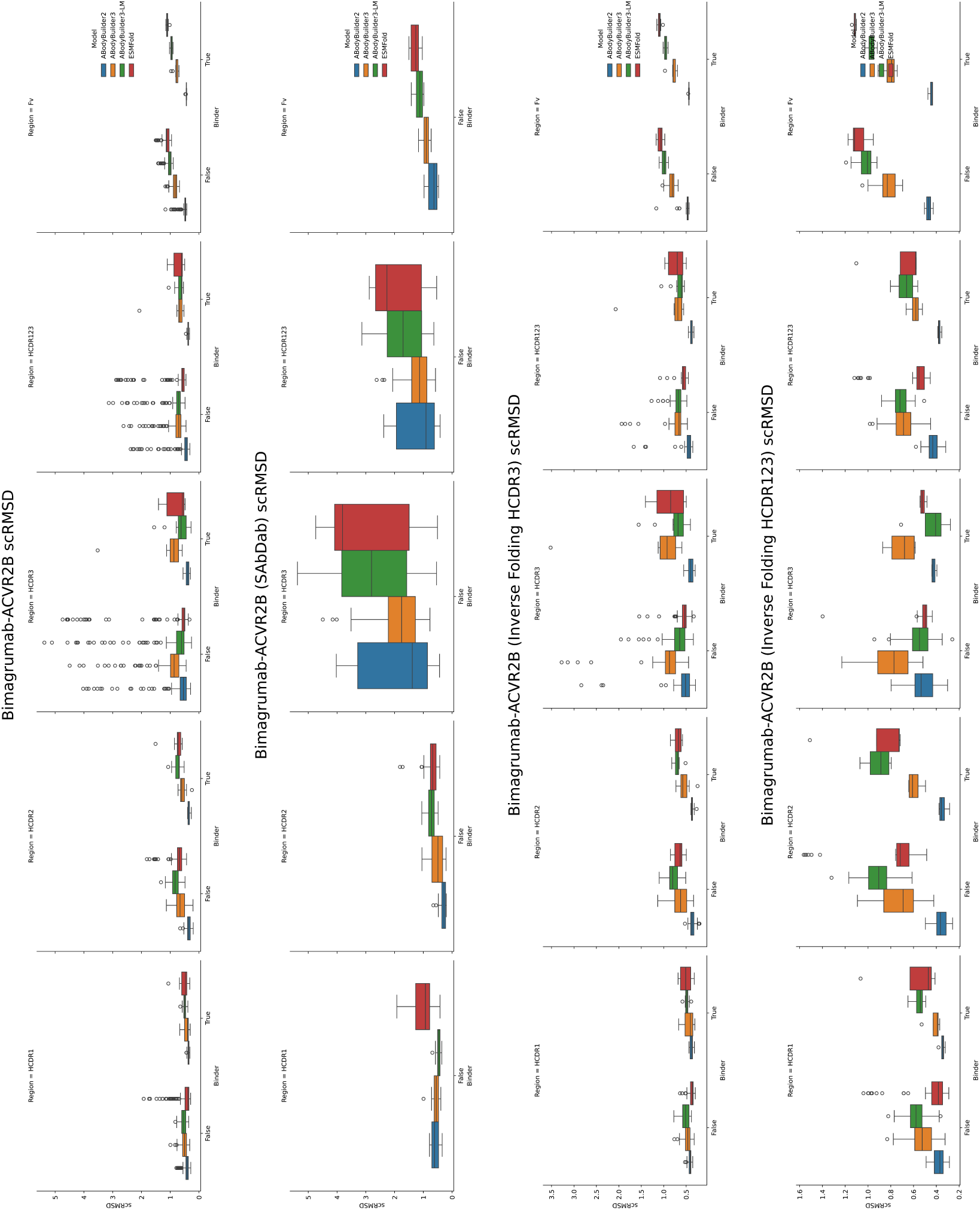
scRMSD distributions of Bimagrumab variants. Values for each of 5 regions (HCDR1, HCDR2, HCDR3, HCDR123, and Fv) and 4 models (ABodyBuilder2, ABodyBuilder3, ABodyBuilder3-LM, and ESMFold) are shown. Distributions are also broken down by binary binding labels and shown (from top to bottom) for all sequences, sequences from the SAbDab baseline, inverse folding HCDR3 designs, and inverse folding HCDR123 designs.

**Figure 22:**
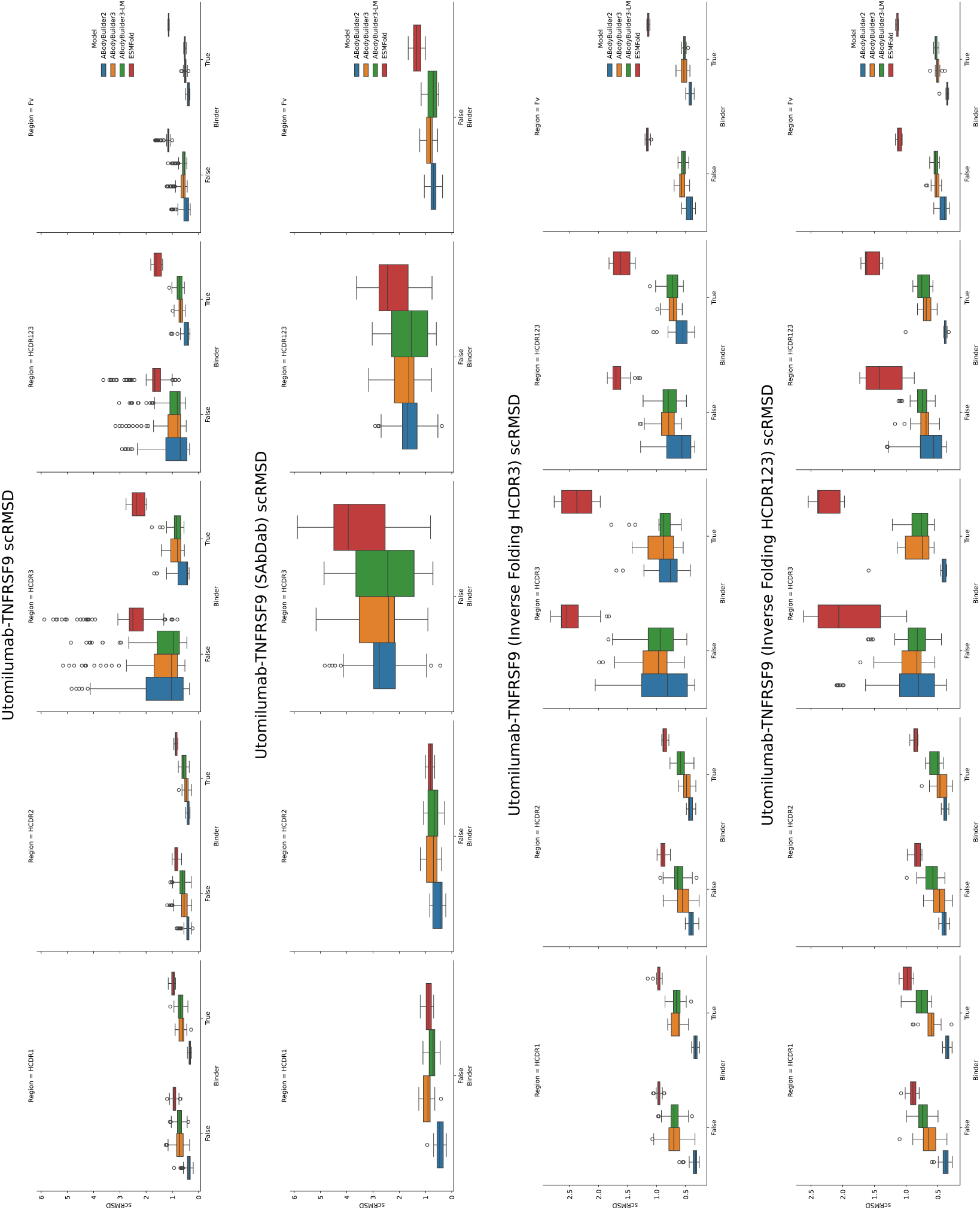
scRMSD distributions of Utomilumab variants. Values for each of 5 regions (HCDR1, HCDR2, HCDR3, HCDR123, and Fv) and 4 models (ABodyBuilder2, ABodyBuilder3, ABodyBuilder3-LM, and ESMFold) are shown. Distributions are also broken down by binary binding labels and shown (from top to bottom) for all sequences, sequences from the SAbDab baseline, inverse folding HCDR3 designs, and inverse folding HCDR123 designs.

We also measured Pearson correlations between binding affinities and scRMSD values as shown in Tables 7, 10, 13, 16, 19, 22, 25, 28. A positive correlation suggests that lower scRMSD corresponds to lower *K_D_* values (higher affinities). For all but one given metrics, that is a folding model and scRMSD over a particular region, we find that correlations can be both positive and negative, suggesting a limited ability to predict affinity. For one metric, Mean Ensemble Fv scRMSD, the correlation is always positive but is as low as 0.001 on Ravagalimab. The median, mean, and standard deviation of correlations for this metric are 0.13, 0.23, and 0.27, The maximum correlation for this metric is 0.87 on Afasevikumab but this is with *n* = 6 binders only.

**Table 19:**
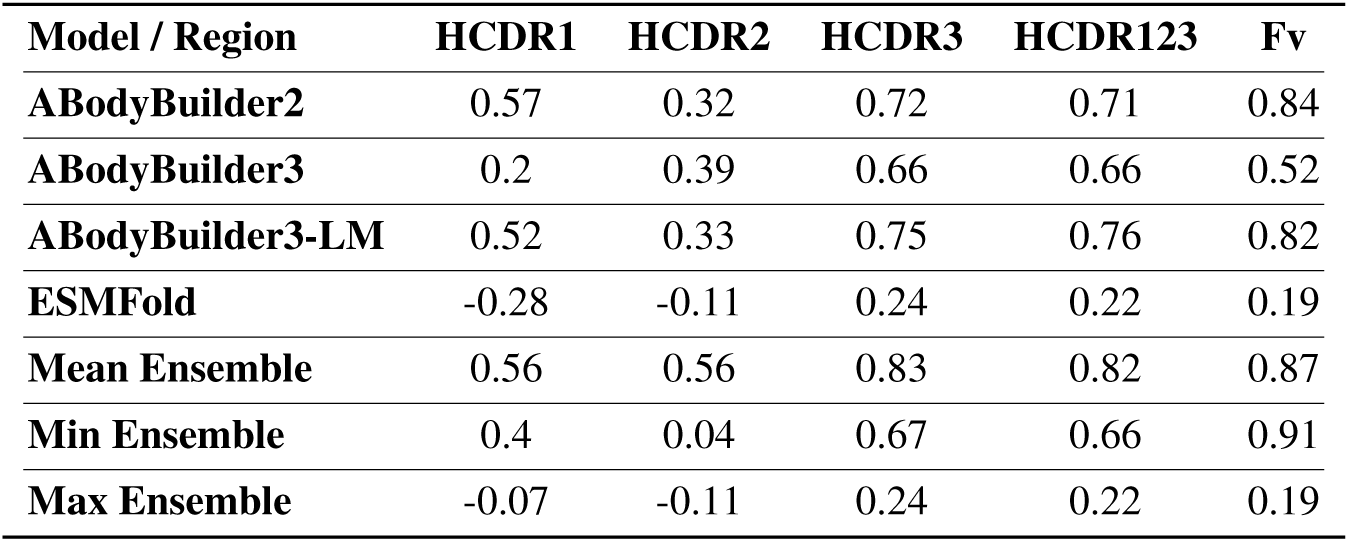
Pearson correlation (*n* = 6) between scRMSD values and affinities (*K_D_* (nM)) of Afasevikumab variant binders to IL17A. A positive correlation implies lower scRMSD values correspond to lower *K_D_* values (higher affinities).

| Model / Region | HCDR1 | HCDR2 | HCDR3 | HCDR123 | Fv |
| --- | --- | --- | --- | --- | --- |
| **ABodyBuilder2** | 0.57 | 0.32 | 0.72 | 0.71 | 0.84 |
| **ABodyBuilder3** | 0.2 | 0.39 | 0.66 | 0.66 | 0.52 |
| **ABodyBuilder3-LM** | 0.52 | 0.33 | 0.75 | 0.76 | 0.82 |
| **ESMFold** | -0.28 | -0.11 | 0.24 | 0.22 | 0.19 |
| **Mean Ensemble** | 0.56 | 0.56 | 0.83 | 0.82 | 0.87 |
| **Min Ensemble** | 0.4 | 0.04 | 0.67 | 0.66 | 0.91 |
| **Max Ensemble** | -0.07 | -0.11 | 0.24 | 0.22 | 0.19 |

**Table 20:** scRMSD values of Osocimab (positive control). Predictions are generated using the unmodified antibody sequence and RMSDs are computed using the solved antibody structure.

| Model / Region | HCDR1 | HCDR2 | HCDR3 | HCDR123 | Fv |
| --- | --- | --- | --- | --- | --- |
| **ABodyBuilder2** | 0.3 | 0.44 | 1.93 | 1.32 | 0.58 |
| **ABodyBuilder3** | 0.52 | 0.52 | 2.16 | 1.5 | 0.67 |
| **ABodyBuilder3-LM** | 0.4 | 0.37 | 1.91 | 1.31 | 0.62 |
| **ESMFold** | 0.77 | 1.1 | 3.52 | 2.46 | 1.13 |

**Table 21:** *t*-test values (*t, p*) between scRMSD values of Osocimab variants and binary binding labels to FXI. A negative *t*-statistic implies mean scRMSD values are lower for binders than non-binders.

| Model / Region | HCDR1 | HCDR2 | HCDR3 | HCDR123 | Fv |
| --- | --- | --- | --- | --- | --- |
| **ABodyBuilder2** | (-0.79, 0.4317) | (1.27, 0.2056) | (-1.71, 0.0904) | (-1.68, 0.0961) | (-1.72, 0.0875) |
| **ABodyBuilder3** | (1.43, 0.1563) | (0.32, 0.7484) | (-0.63, 0.5299) | (-0.53, 0.5969) | (-0.23, 0.8207) |
| **ABodyBuilder3-LM** | (-3.24, 0.0016) | (-2.2, 0.0296) | (-1.55, 0.1246) | (-1.66, 0.0993) | (-1.67, 0.0968) |
| **ESMFold** | (-1.85, 0.0676) | (-0.02, 0.9855) | (-2.02, 0.0459) | (-2.02, 0.0459) | (-2.24, 0.0273) |
| **Mean Ensemble** | (-1.77, 0.0789) | (-0.09, 0.929) | (-2.08, 0.0394) | (-2.09, 0.0384) | (-2.14, 0.0343) |
| **Min Ensemble** | (-1.31, 0.1931) | (-1.61, 0.1112) | (-1.16, 0.2495) | (-0.7, 0.4852) | (-0.53, 0.5967) |
| **Max Ensemble** | (-1.69, 0.093) | (-0.63, 0.5273) | (-2.71, 0.0077) | (-2.81, 0.0058) | (-2.59, 0.0109) |

**Table 22:** Pearson correlation (*n* = 34) between scRMSD values and affinities (*K_D_* (nM)) of Osocimab variant binders to FXI. A positive correlation implies lower scRMSD values correspond to lower *K_D_* values (higher affinities).

| Model / Region | HCDR1 | HCDR2 | HCDR3 | HCDR123 | Fv |
| --- | --- | --- | --- | --- | --- |
| **ABodyBuilder2** | -0.14 | -0.16 | 0.26 | 0.25 | 0.22 |
| **ABodyBuilder3** | 0.0 | 0.13 | -0.21 | -0.2 | -0.07 |
| **ABodyBuilder3-LM** | -0.11 | 0.15 | 0.25 | 0.26 | 0.25 |
| **ESMFold** | -0.18 | -0.17 | -0.09 | -0.12 | -0.12 |
| **Mean Ensemble** | -0.17 | -0.03 | 0.07 | 0.08 | 0.13 |
| **Min Ensemble** | -0.07 | 0.12 | -0.24 | -0.21 | 0.14 |
| **Max Ensemble** | -0.24 | -0.16 | 0.17 | 0.15 | -0.06 |

**Table 23:**
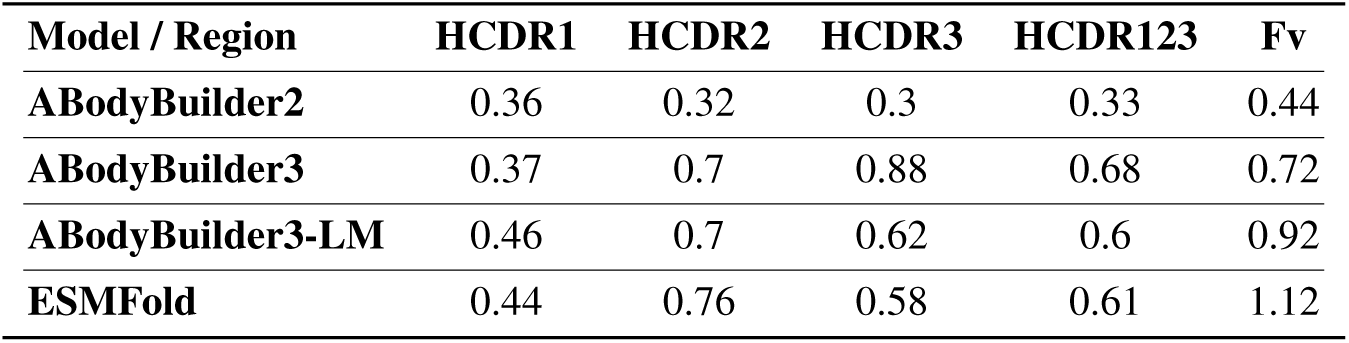
scRMSD values of Bimagrumab (positive control). Predictions are generated using the unmodified antibody sequence and RMSDs are computed using the solved antibody structure.

| Model / Region | HCDR1 | HCDR2 | HCDR3 | HCDR123 | Fv |
| --- | --- | --- | --- | --- | --- |
| **ABodyBuilder2** | 0.36 | 0.32 | 0.3 | 0.33 | 0.44 |
| **ABodyBuilder3** | 0.37 | 0.7 | 0.88 | 0.68 | 0.72 |
| **ABodyBuilder3-LM** | 0.46 | 0.7 | 0.62 | 0.6 | 0.92 |
| **ESMFold** | 0.44 | 0.76 | 0.58 | 0.61 | 1.12 |

**Table 24:** *t*-test values (*t, p*) between scRMSD values of Bimagrumab variants and binary binding labels to ACVR2B. A negative *t*-statistic implies mean scRMSD values are lower for binders than non-binders.

| Model / Region | HCDR1 | HCDR2 | HCDR3 | HCDR123 | Fv |
| --- | --- | --- | --- | --- | --- |
| **ABodyBuilder2** | (-2.22, 0.0279) | (0.11, 0.9117) | (-1.84, 0.0674) | (-1.79, 0.0758) | (-2.14, 0.0338) |
| **ABodyBuilder3** | (-2.47, 0.0145) | (-1.83, 0.0683) | (-0.21, 0.8317) | (-0.67, 0.5027) | (-1.97, 0.0504) |
| **ABodyBuilder3-LM** | (-1.16, 0.2464) | (-1.79, 0.0757) | (-1.12, 0.2663) | (-1.47, 0.1432) | (-2.61, 0.0098) |
| **ESMFold** | (0.16, 0.8708) | (0.45, 0.6567) | (-0.53, 0.5945) | (-0.46, 0.6456) | (-0.45, 0.6534) |
| **Mean Ensemble** | (-1.28, 0.2021) | (-1.28, 0.2014) | (-1.02, 0.3099) | (-1.2, 0.2336) | (-2.31, 0.0219) |
| **Min Ensemble** | (-1.07, 0.2869) | (-0.17, 0.8629) | (-1.71, 0.0891) | (-1.97, 0.0504) | (-2.28, 0.024) |
| **Max Ensemble** | (-0.81, 0.4195) | (-1.22, 0.2244) | (-0.49, 0.623) | (-0.64, 0.5257) | (-0.76, 0.4492) |

**Table 25:** Pearson correlation (*n* = 14) between scRMSD values and affinities (*K_D_* (nM)) of Bimagrumab variant binders to ACVR2B. A positive correlation implies lower scRMSD values correspond to lower *K_D_* values (higher affinities).

| Model / Region | HCDR1 | HCDR2 | HCDR3 | HCDR123 | Fv |
| --- | --- | --- | --- | --- | --- |
| **ABodyBuilder2** | -0.23 | -0.41 | 0.51 | 0.14 | 0.09 |
| **ABodyBuilder3** | -0.05 | 0.23 | -0.33 | -0.29 | 0.15 |
| **ABodyBuilder3-LM** | -0.09 | 0.38 | -0.2 | -0.03 | 0.16 |
| **ESMFold** | 0.26 | 0.22 | -0.06 | 0.17 | -0.13 |
| **Mean Ensemble** | 0.13 | 0.36 | -0.26 | -0.15 | 0.14 |
| **Min Ensemble** | -0.19 | -0.18 | 0.21 | 0.14 | 0.09 |
| **Max Ensemble** | 0.25 | 0.38 | -0.2 | -0.02 | -0.13 |

**Table 26:**
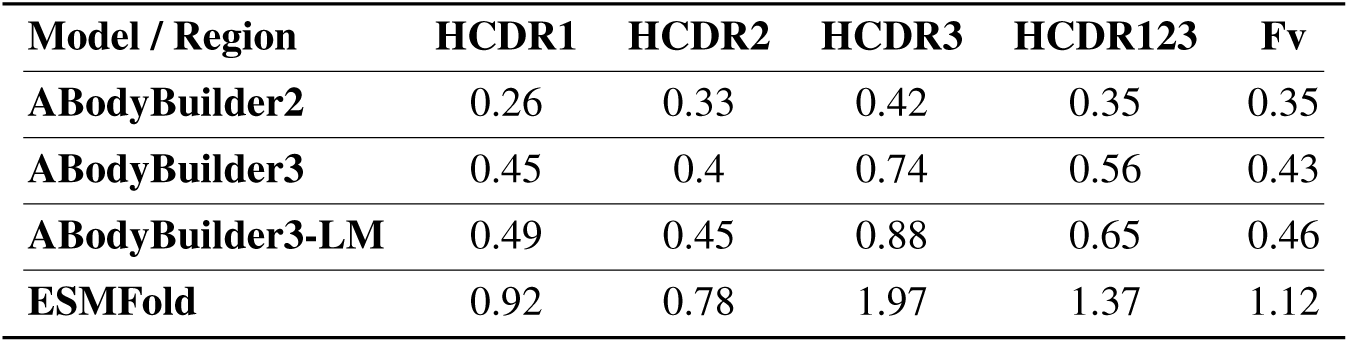
scRMSD values of Utomilumab (positive control). Predictions are generated using the unmodified antibody sequence and RMSDs are computed using the solved antibody structure.

| Model / Region | HCDR1 | HCDR2 | HCDR3 | HCDR123 | Fv |
| --- | --- | --- | --- | --- | --- |
| **ABodyBuilder2** | 0.26 | 0.33 | 0.42 | 0.35 | 0.35 |
| **ABodyBuilder3** | 0.45 | 0.4 | 0.74 | 0.56 | 0.43 |
| **ABodyBuilder3-LM** | 0.49 | 0.45 | 0.88 | 0.65 | 0.46 |
| **ESMFold** | 0.92 | 0.78 | 1.97 | 1.37 | 1.12 |

**Table 27:**
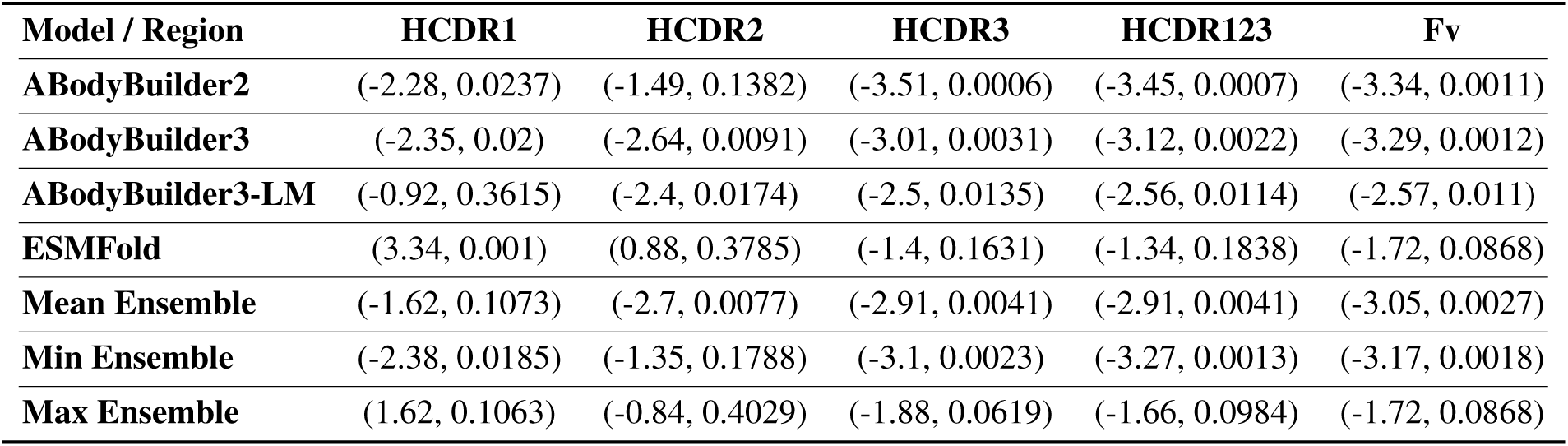
*t*-test values (*t, p*) between scRMSD values of Utomilumab variants and binary binding labels to TNFRSF9. A negative *t*-statistic implies mean scRMSD values are lower for binders than non-binders.

**Table 28:** Pearson correlation (*n* = 28) between scRMSD values and affinities (*K_D_* (nM)) of Utomilumab variant binders to TNFRSF9. A positive correlation implies lower scRMSD values correspond to lower *K_D_* values (higher affinities).

| Model / Region | HCDR1 | HCDR2 | HCDR3 | HCDR123 | Fv |
| --- | --- | --- | --- | --- | --- |
| **ABodyBuilder2** | -0.01 | 0.05 | 0.29 | 0.27 | 0.18 |
| **ABodyBuilder3** | -0.12 | -0.29 | 0.0 | -0.09 | 0.19 |
| **ABodyBuilder3-LM** | 0.09 | 0.05 | 0.12 | 0.13 | -0.05 |
| **ESMFold** | 0.02 | 0.14 | 0.22 | 0.21 | 0.06 |
| **Mean Ensemble** | -0.02 | -0.11 | 0.28 | 0.26 | 0.2 |
| **Min Ensemble** | -0.12 | -0.09 | 0.34 | 0.3 | 0.18 |
| **Max Ensemble** | 0.0 | 0.14 | 0.22 | 0.21 | 0.06 |

## 4 Discussion

Here we have presented IgDesign, an antibody inverse folding model developed by combining (1) ideas from protein inverse folding models and language models such as ProteinMPNN, LM-Design, and ESM2, (2) an antibody-specific framing of the problem with antigen and antibody FWR sequences provided as context, and (3) fine-tuning on antibody-antigen complexes. We demonstrate IgDesign’s ability to consistently design binders against multiple target antigens with confirmation using SPR.

Using the SPR datasets generated by evaluating IgDesign, we evaluate the performance of self-consistency RMSD (scRMSD) in discriminating between binders and non-binders as well as for predicting binding affinity. We find limited evidence of the usefulness of this metric, motivating a more explicit evaluation of scRMSD as well as the development of different metrics for antibody design tasks. We open source these datasets to facilitate future benchmarking efforts by the community.^7^

We present the first *in vitro* validation of inverse folding for antibody binder design against a diverse set of therapeutic antigens. Demonstrating the success of antibody inverse folding is key to advancing the field since models such as IgDesign can be broadly applied to antibody development efforts. In particular, IgDesign can enhance lead optimization efforts, for example with superior variant prediction, and enable *de novo* antibody design consisting of structure design followed by inverse folding.

## Acknowledgments

The authors wish to thank Janani S. Iyer, Simon V. Mathis, Kieran Didi, Amaro Taylor-Weiner, Byron Olsen, Kristin Iannacone, and Shelby Walker for critical review of this manuscript; Daniele Biasci, Miles Gander, Jens Plassmeier, and James Mategko for early discussion and planning of this effort; Zach Jonasson, Andreas Busch, and Sean McClain for continual support.

## A Modeling Methods

### A.1 IgMPNN

IgMPNN takes as input the 3Dcoordinates of the backbone residues of an antibody-antigen complex. We define a protein graph as a directed graph where residues are represented as nodes and share edges with their *k* nearest neighbors. We use *k* = 48 in all of our experiments. We initialize the node features *x_i_* and edge features *e^t^* in our protein graph using the following features from [1]: (1) Distances between C*α*-C*α* atoms, (2) Relative C*α*-C*α*-C*α* frame orientations and rotations, (3) Backbone dihedral angles, (4) Binary features that determine relative chain positions, and (5) Relative position encodings. Our featurization differs from [1]in two ways: (1) We do not assume access to any side chain atoms and thus we do not include any pairwise distance features involving side chain atoms. (2) We include embedded residue type features for all antigen residues and antibody framework residues. We replace the antibody CDR residue embeddings with zero vectors.

Our initial features, *x*_0_, then get passed into our message passing neural network encoder. Our network has multiple message passing phases during which the hidden state of each node in the graph *h^t^* and edge *e^t^* is updated according to:

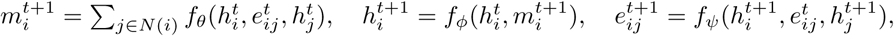

where *f_θ_* is our message update function, *f_ϕ_* is our node update function, *f_ψ_* is our edge update function, and *N* (*i*) is the set of neighboring nodes for a given node *i* in the graph. We use 128 as our hidden node dimension throughout the network. IgMPNN utilizes three encoder layers, performing message passing node updates and edge updates. The output is fed into the decoder, which has three layers. The decoder performs message passing to update the node representations according 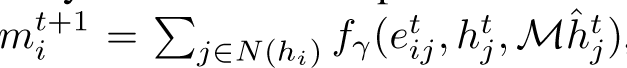, where *f_γ_* is our decoder message update function. The ground truth context, *h*^^*t*^, which is an embedding of the ground truth residues, is provided as input only when it is allowed by our causal decoding mask, *M*. The decoder is masked to prevent the model from incorporating information from nodes that have yet to be decoded while allowing the decoder to access information from nodes that have already been decoded. When decoding a given residue during training time, instead of accessing the embeddings for predicted residues at previously decoded positions, the decoder accesses embeddings for ground truth residues at previously decoded positions. The model decodes antibody CDRs in sequential order during training: HCDR1, HCDR2, HCDR3, LCDR1, LCDR2, LCDR3. During inference, a different order of CDRs can be specified. Within each CDR, the decoding order is random as done in [1]. We project the final node embeddings to logits and train the model using cross-entropy loss.

### A.2 Discussion of ESM and potential data leakage

Data leakage from ESM [13, 14] is a concern that was originally raised for general protein inverse folding with LM-Design [3]. The authors presented an argument against this concern^8^. It has also been noted that the number of antibody-related sequences ESM has been trained on is in the thousands compared to the multiple tens of millions of proteins in its training set [26], which suggests it is less likely to observe leakage for antibody design compared to general protein design. Furthermore, our *in silico* results show that IgMPNN, which does not use ESM, achieves comparable HCDR AARs to IgDesign, implying that information is not being leaked from ESM. Finally, we consider data leakage for ProteinMPNN [1], noting in our *in silico* results that despite training on >80% of the antibody-antigen complexes in the relevant test sets, ProteinMPNN still underperforms both IgMPNN and IgDesign. While ProteinMPNN and ESM are considerably different models, this example motivates the fact that models can ignore elements of their training sets.

### A.3 ProteinMPNN *in silico* baseline

We perform an *in silico* baseline against ProteinMPNN by taking the open source implementation of the model^9^ and running inference on the antibody-antigen complexes in the test sets of the IgDesign models we trained. Inference is run with default parameters directly from the PDB files^10^. We generate one sequence for each complex. We then parse the CDRs from these sequences and compute 1-shot AARs.

We investigated the training set of ProteinMPNN and noted that a significant portion of SAbDab [17] is in the training set. Indeed, nearly 25,000 non-unique PDB IDs from SAbDab are contained in ProteinMPNN’s training set. The model is primarily trained on individual chains from these complexes (i.e., heavy chains, light chains, or antigens treated as monomers). This results in *≈*80% of our test set complexes being contained in ProteinMPNN’s training set. Because of this, we also compute the “ProteinMPNN (Filtered)” baseline by restricting to antibodies contained in IgDesign’s test sets that are not contained in ProteinMPNN’s training set as monomers (i.e., if the heavy or light chain appears in ProteinMPNN’s training set as a monomer it will not be considered in this baseline).

Interestingly, despite this data leakage, IgDesign outperforms ProteinMPNN with statistical significance on every CDR. Furthermore, ProteinMPNN and ProteinMPNN (Filtered) perform comparably to each other.

### A.4 Self-consistency RMSD (scRMSD) computation

ABodyBuilder2 [6], ABodyBuilder3 [7], and ESMFold [8] were used to predict Fv structures of all antibody heavy and light chain variable sequences. For ABodyBuilder3 we also generated predictions with ABodyBuilder3 using ProtT5 [27] for generating embeddings. For ESMFold the heavy and light chain were concatenated with a linker consisting of 25 Glycine residues in between them. After structure prediction, the linker was removed and the heavy and light predictions were treated as separate chains.

Model predicted Fv structures were Kabsch-aligned on C*α* residues to the structure of the reference antibody. PDBs and chain IDs used for reference antibodies can be found in Table 2. After alignment, C*α* RMSDs were computed on HCDR1, HCDR2, HCDR3, HCDR123, and the entire Fv. In the event of missing FWR residues in the PDB structures, these residues were removed from consideration during alignemnt and RMSD calculation. We refer to these RMSD values as the self-consistency RMSD or scRMSD. Note that scRMSD is not only a function of the antibody sequence and its reference antibody structure, but also of the model used for folding the sequence and the region under consideration.

## B Data curation and splitting

The Structural Antibody Database (SAbDab) [17] was retrieved on December 6th, 2022. The corresponding PDB files were downloaded from RSCB PDB [22]. To generate a high quality dataset used in training we applied the following filters:

- Drop entries without a heavy antibody chain.
- Drop entries without an antigen.
- Drop entries where the PDB id, heavy, light, and antigen chains are repeated (duplicated).
- Drop entries where one of the heavy CDRs is too short (shorter than 5 amino acids for HCDR1 and HCDR2, shorter than 7 amino acids for HCDR3).
- Drop entries where one of the heavy CDRs is too long (longer than 26 amino acids for any of the HCDRs).
- Drop entries where more than 10% of heavy chain residues are missing from the structure.
- Drop entries where more than 25% of antigen residues are missing from the structure.

After filtering there were 6933entries left in the database. To split the dataset, we use a 40% antigen sequence identity holdout. Specifically, we applied sequence clustering to the antigen sequences using mmseqs2 [28] version 13.45111, with the following parameters: min-seq-id=0.4, cov-mode=1, cluster-mode=2, cluster-reassign=true. We use these cluster annotations when splitting the data into train, validation, and test folds (assigning an entire cluster to one of the three subsets).

The General PDB dataset is created from a selection of PDB files available in the RSCB PDB [22] database. We made the selection using the Graphein library (version 1.0) [29], downloading all PDB structures where each chain is longer than 40 and shorter than 500 amino acids. We further filtered out any structures containing chains with missing backbone atoms. Finally, we removed all PDBs contained in SAbDab to avoid leakage. This resulted in a dataset with 74734 entries. We then applied sequence clustering using mmseqs2 with the same set of parameters used for clustering the SAbDab dataset (min-seq-id=0.4, cov-mode=1, cluster-mode=2, cluster-reassign=true). As with SAbDab, we use these cluster annotations when splitting the data into train, validation, and test folds (assigning an entire cluster to one of the three subsets).

## C Additional details on antibody library design

We make note of additional details for antibody library design:

- For each of the 8 target antigens we trained IgDesign with a SAbDab split holding out said antigen.
- As mentioned in the main text, we generated sequences in the following order: HCDR3, HCDR1, HCDR2, LCDR1, LCDR2, LCDR3. We selected this order to prevent the model from conditioning on potentially false CDR predictions when predicting HCDR3. We note that the model is never provided ground truth CDR sequences at inference time.
- For generating sequences with IgDesign we used weighted random sampling after applying softmax with a temperature of *T* = 0.5 to the model’s logits.
- As a baseline, we sampled 100 HCDR3s from the training set (a subset of SAbDab) of each IgDesign model. We required these HCDR3s to have the same length as the HCDR3 of the reference antibody. If there were fewer than 100 such HCDR3s, we first took HCDR3s from the validation set with matching lengths then, if needed, we took HCDR3s 1 residue longer or shorter from the training set.
- We included controls for the SPR assay to confirm its ability to properly label known binders and non-binders. In particular, for each experiment we included a monoclonal antibody (mAb) positive control (except for CD40), a fragment antigen-binding (Fab) positive control and, and a Fab negative control.

### Discussion of the *in vitro* baseline

We describe the construction of the baseline above. The objective of the baseline is to compare to IgDesign’s binding rates and demonstrate that the model is relying on its input features (antibody-antigen complex structure, antigen sequence, antibody FWR sequences) and that it has learned to extrapolate from its training set of HCDRs. By sampling HCDR3s from the model’s training set, we can effectively answer these questions. The success rates of the baseline are low. In fact, they are 0% for 6 out of 8 antigens and for the remaining 2 each baseline population contains only 1 binder, which corresponds to a success rate of *≈* 1.5% (Table 3). Given IgDesign’s outperformance vs. the baseline, we can conclude that the model has learned to extrapolate by relying on its input features.

We note that there are multiple other baselines of interest. The SAbDab HCDR3 baseline does not utilize IgDesign’s input features for sampling sequences. As a result, while we can determine that the model uses these features we cannot determine how effectively it uses these features. A baseline using a protein inverse folding model such as ProteinMPNN [1] or an antibody inverse folding model such as AbMPNN [5] would help answer this question. Similarly, baselines using protein language models such as ESM [13, 14] or antibody language models such as IgLM [30] would help assess the importance of the structure features. There are, however, some shortcomings of these baselines compared to our SAbDAb HCDR3 baseline. Firstly, we do not have a guarantee that the CDRs generated by these models are realistic. Indeed, AbMPNN [5] shows that ProteinMPNN’s CDR designs have substantially lower designability and accuracy (as measured by AAR) compared to AbMPNN. We show similar results for AAR with IgDesign outperforming ProteinMPNN. Secondly, as this study demonstrates the first effort at multi-antigen *in vitro* validation of antibody inverse folding, we do not yet have an existing antibody inverse folding model that has been validated to compare against. Having experimentally validated antibody inverse folding, future work can consider additional baselines including the models suggested above as well as both our SAbDAb baseline and our model, IgDesign.

## D Wet lab workflow

### D.1 Cloning

Antibody variants are cloned and expressed in fragment antigen-binding (Fab) format. To produce SPR datasets, DNA variants spanning HCDR1 to HCDR3 are purchased as single-stranded DNA (ssDNA) oligo pools (Twist Bioscience). For each residue, codons are randomly selected uniformly from the codons that translate into said residue.

Amplification of the ssDNA oligo pools is carried out by PCR according to Twist Bioscience’s recommendations, except Q5 high fidelity DNA polymerase (New England Biolabs) is used in place of KAPA polymerase. Briefly, 25 *µ*L reactions consist of 1x Q5 Mastermix, 0.3 *µ*M each of forward and reverse primers, and 10 ng oligo pool. Reactions are initially denatured for 3 min at 95^◦^C, followed by 13 cycles of: 95^◦^C for 20 s; 66^◦^C for 20 s; 72^◦^C for 15 s; and a final extension of 72^◦^C for 1 min. DNA amplification is confirmed by agarose gel electrophoresis, and amplified DNA is subsequently purified (DNA Clean and Concentrate Kit, Zymo Research).

To generate linearized vector, a two-step PCR is carried out to split our plasmid vector carrying Fab format antibody into two fragments in a manner that provides cloning overlaps of approximately 25 nucleotides (nt) on the 5’ and 3’ ends of the amplified ssDNA oligo pool libraries, or 40 nt on the 5’ and 3’ ends of IDT eBlocks. Vector linearization reactions are digested with DpnI (New England Bioloabs) and purified from a 0.8% agarose gel using the Gel DNA Recovery Kit (Zymo Research) to eliminate parental vector carry through. Cloning reactions consist of 50 fmol of each purified vector fragment, either 100 fmol PCR-amplified ssDNA oligo pool or 10 pmol eBlock library inserts and 1x final concentration NEBuilder HiFi DNA Assembly (New England Biolabs). Reactions are incubated at 50^◦^C for 25 min using eBlocks or two hours using PCR-amplified oligo pools. Assemblies are subsequently purified using the DNA Clean and Concentrate Kit (Zymo Research). DNA concentrations are measured using a NanoDrop OneC (Thermo Scientific).

For SPR datasets, the *E. coli* host strain is transformed with the purified assembly reactions and grown overnight at 30^◦^C on agar plates containing 50 *µ*g/ml kanamycin and 1 % glucose. Colonies are picked for QC analysis prior to cultivation for induction.

#### QC Analysis

Quality of antibody variant libraries is assessed by performing rolling circle amplification (Equiphi29, Thermo Fisher Scientific) on 24 colonies and sequencing using the Illumina DNA Prep, Tagmentation Kit (Illumina Inc.). Each colony is analyzed for mutations from reference sequence, presence of multiple variants, misassembly, and matching to a library sequence (Geneious Prime).

### D.2 Antibody Expression in *E. coli*

After transformation and QC of libraries, individual colonies are picked into deep well plates containing 400 *µ*L of Teknova LB Broth 50 *µ*g/mL Kanamycin and incubated at 30^◦^C and 80 % humidity with 1000 rpm shaking for 24 hours. At the end of the 24 hours, 150 *µ*L samples are centrifuged (3300 g, 7 min), supernatant decanted from the pre-culture plate, and cell pellets sent for sequence analysis. 80 *µ*L of the pre-culture is transferred to 400 *µ*L of IBM containing inducers and supplements as described above. Culture is grown for 16 hours at 26^◦^C and 80 % humidity with 270 rpm shaking. After 16 hours, 150 *µ*L samples are taken and centrifuged (3300 g, 7 min) into pellets with supernatant decanting prior to being stored at -80^◦^C.

### D.3 Surface plasmon resonance (SPR)

#### Sample Preparation

Post induction samples are transferred to 96-well plates (Greiner Bio-One), pelleted and lysed in 50 *µ*L lysis buffer (1X BugBuster protein extraction reagent containing 0.01 KU Benzonase Nuclease and 1X Protease inhibitor cocktail). Plates are incubated for 15-20 min at 30^◦^C then centrifuged to remove insoluble debris. After lysis, samples are adjusted with 200 *µ*L SPR running buffer (10 mM HEPES, 150 mM NaCl, 3 mM EDTA, 0.01 % w/v Tween-20, 0.5 mg/mL BSA) to a final volume of 260 *µ*L and filtered into 96-well plates. Lysed samples are then transferred from 96-well plates to 384-well plates for high-throughput SPR using a Hamilton STAR automated liquid handler. Colonies are prepared in two sets of independent replicates prior to lysis and each replicate is measured in two separate experimental runs. In some instances, single replicates are used, as indicated.

#### SPR

High-throughput SPR experiments are conducted on a microfluidic Carterra LSA SPR instrument using SPR running buffer (10 mM HEPES, 150 mM NaCl, 3 mM EDTA, 0.01 % w/v Tween-20, 0.5 mg/mL BSA) and SPR wash buffer (10 mM HEPES, 150 mM NaCl, 3 mM EDTA, 0.01 % w/v Tween-20). Carterra LSA SAD200M chips are pre-functionalized with 20 *µ*g/mL biotinylated antibody capture reagent for 600 s prior to conducting experiments. Lysed samples in 384-well blocks are immobilized onto chip surfaces for 600 s followed by a 60 s washout step for baseline stabilization. Antigen binding is conducted using the non-regeneration kinetics method with a 300 s association phase followed by a 900 s dissociation phase. For analyte injections, six leading blanks are introduced to create a consistent baseline prior to monitoring antigen binding kinetics. After the leading blanks, five concentrations of antigen extracellular domain antigen (ACRO Biosystems, prepared in three-fold serial dilution from a starting concentration of 500 nM), are injected into the instrument and the time series response was recorded. In most experiments, measurements on individual DNA variants are repeated four times. Typically each experiment run consists of two complete measurement cycles (ligand immobilization, leading blank injections, analyte injections, chip regeneration) which provide two duplicate measurement attempts per clone per run. In most experiments, technical replicates measured in separate runs further double the number of measurement attempts per clone to four.

### D.4 Sequencing

To identify the DNA sequence of individual antibody variants evaluated by SPR, duplicate plates are provided for sequencing. A portion of the pelleted material is transferred into 96 well PCR (Thermo-Fisher) plate via pinner (Fisher Scientific) which contains reagents for performing an initial phase PCR of a two-phase PCR for addition of Illumina adapters and sequencing. Reaction volumes used are 12.5 *µ*l. During the initial PCR phase, partial Illumina adapters are added to the amplicon via 4 PCR cycles. The second phase PCR adds the remaining portion of the Illumina sequencing adapter and the Illumina i5 and i7 sample indices. The initial PCR reaction uses 0.45 *µ*M UMI primer concentration, 6.25 *µ*l Q5 2x master mix (New England Biolabs) and PCR grade H_2_O. Reactions are initially denatured at 98^◦^C for 3 min, followed by 4 cycles of 98^◦^C for 10 s; 59^◦^C for 30 s; 72^◦^C for 30 s; with a final extension of 72^◦^C for 2 min. Following the initial PCR, 0.5 *µ*M of the secondary sample index primers are added to each reaction tube. Reactions are then denatured at 98^◦^C for 3 min, followed by 29 cycles of 98^◦^C for 10 s; 62^◦^C for 30 s; 72^◦^C for 15 s; with a final extension of 72^◦^C for 2 min. Reactions are then pooled into a 1.5 mL tube (Eppendorf). Pooled samples are size selected with a 1x AMPure XP (Beckman Coulter) bead procedure. Resulting DNA samples are quantified by Qubit fluorometer. Pool size is verified via Tapestation 1000 HS and is sequenced on an Illumina MiSeq Reagent Kit v3 (2×300 nt) for HCDR1-HCDR3 libraries with 20 % PhiX.

After sequencing, amplicon reads are merged using Fastp [31], trimmed by cutadapt [32] and each unique sequence enumerated. Next, custom R scripts are applied to calculate sequence frequency ratios between the most abundant and second-most abundant sequence in each sample. Levenshtein distance is also calculated between the two sequences. These values are used for downstream filtering to ensure a clonal population is measured by SPR. The most abundant sequence within each sample is compared to the designed sequences and discarded if it does not match any expected sequence. Dominant sequences are then combined with their companion Carterra SPR measurements.

## E Additional figures and tables

### E.1 Amino acid recovery

### E.2 *In vitro* binding rates

### E.3 *In vitro* binding affinities

We show binding affinities for binders against each antigen from IgDesign as well as the SAbDab HCDR3 baseline. The reference antibody affinity is shown in each figure by a dashed black line. For affinities we report *−* log_10_(*K_D_*) (*M*).

For 5 out of 8 antigens (CD40, IL36R, FXI, ACVR2B, TNFRSF9), IgDesign generates binders with equal or higher affinities to the reference antibody. For these 5 targets, several designed binders have affinities within one order of magnitude of the reference antibody’s affinity. This suggests that IgDesign can be used for lead optimization, in particular affinity maturation, either by designing HCDRs or via an efficient evolution approach [26, 19] by ranking HCDR mutants conditional on a backbone structure. For the other 3 out of 8 antigens (C5, TSLP, IL17A), the binders are 2+ orders of magnitude lower affinity than reference, motivating future work to improve the model.

### E.4 Assessment of self-consistency RMSD (scRMSD)

We show scRMSD distributions and statistics for all evaluated sequences across the 8 antigens considered. Figures showing scRMSD distributions are shown for each antigen in Figures 15,16,17,18,19,20,21, 22. Reference scRMSD values, this is the scRMSD of the reference antibody sequence itself, are shown in Tables 5, 8, 11, 14, 17, 20, 23, 26. Two-sided *t*-tests on scRMSD between binders and non-binders are shown in Tables 6, 9, 12, 15, 18, 21, 24, 27. Pearson correlations between scRMSD and affinities are shown in Tables 7, 10, 13, 16, 19, 22, 25, 28.

## Footnotes

2 The term *reference antibody* refers to the antibody in the antibody-antigen complex used for inverse folding.

3 Code and datasets can be found at https://github.com/AbSciBio/igdesign. At this time, we have released datasets for 7 out of 8 antibody-antigen systems used to evaluate IgDesign.

4 We make an argument for why potential data leakage from ESM is not a concern in the Appendix.

5 We include details on the ProteinMPNN *in silico* baseline in the Appendix.

6 We require *p* < 3e-3 for statistical significance as we are considering *α* = 0.05 with *N* = 16 tests.

7 Code and datasets can be found at https://github.com/AbSciBio/igdesign.

8 https://github.com/BytedProtein/ByProt/issues/3

9 https://github.com/dauparas/ProteinMPNN

10 https://github.com/dauparas/ProteinMPNN/blob/main/examples/submit_example_3.sh

## References

[1] J. Dauparas, I. Anishchenko, N. Bennett, H. Bai, R. J. Ragotte, L. F. Milles, B. I. M. Wicky, A. Courbet, R. J. de Haas, N. Bethel, P. J. Y. Leung, T. F. Huddy, S. Pellock, D. Tischer, F. Chan, B. Koepnick, H. Nguyen, A. Kang, B. Sankaran, A. K. Bera, N. P. King, and D. Baker. Robust deep learning–based protein sequence design using ProteinMPNN. Science, 378(6615):49–56, October 2022. doi: 10.1126/science.add2187. URL 10.1126/science.add2187.

[2] Chloe Hsu, Robert Verkuil, Jason Liu, Zeming Lin, Brian Hie, Tom Sercu, Adam Lerer, and Alexander Rives. Learning inverse folding from millions of predicted structures. In Kamalika Chaudhuri, Stefanie Jegelka, Le Song, Csaba Szepesvari, Gang Niu, and Sivan Sabato, editors, Proceedings of the 39th International Conference on Machine Learning, volume 162 of Proceedings of Machine Learning Research, pages 8946–8970. PMLR, 17–23 Jul 2022. URL https://proceedings.mlr.press/v162/hsu22a. html.

[3] Zaixiang Zheng, Yifan Deng, Dongyu Xue, Yi Zhou, Fei YE, and Quanquan Gu. Structure-informed language models are protein designers, 2023.

[4] Sai Pooja Mahajan, Jeffrey A. Ruffolo, and Jeffrey J. Gray. Contextual protein and antibody encodings from equivariant graph transformers. July 2023. doi: 10.1101/2023.07.15.549154. URL https://doi.org/10.1101/2023.07.15.549154.

[5] Frédéric A Dreyer, Daniel Cutting, Constantin Schneider, Henry Kenlay, and Charlotte M Deane. Inverse folding for antibody sequence design using deep learning. ICML Computational Biology Workshop Poster, 2023.

[6] Brennan Abanades, Wing Ki Wong, Fergus Boyles, Guy Georges, Alexander Bujotzek, and Charlotte M. Deane. Immunebuilder: Deep-learning models for predicting the structures of immune proteins. Communications Biology, 6(1), May 2023. ISSN 2399-3642. doi: 10.1038/s42003-023-04927-7. URL 10.1038/s42003-023-04927-7.

[7] Henry Kenlay, Frédéric A. Dreyer, Daniel Cutting, Daniel Nissley, and Charlotte M. Deane. Abodybuilder3: Improved and scalable antibody structure predictions, 2024.

[8] Zeming Lin, Halil Akin, Roshan Rao, Brian Hie, Zhongkai Zhu, Wenting Lu, Nikita Smetanin, Robert Verkuil, Ori Kabeli, Yaniv Shmueli, Allan dos Santos Costa, Maryam Fazel-Zarandi, Tom Sercu, Salvatore Candido, and Alexander Rives. Evolutionary-scale prediction of atomic-level protein structure with a language model. Science, 379(6637):1123–1130, March 2023. ISSN 1095-9203. doi: 10.1126/science.ade2574. URL 10.1126/science.ade2574.

[9] Joseph L. Watson, David Juergens, Nathaniel R. Bennett, Brian L. Trippe, Jason Yim, Helen E. Eisenach, Woody Ahern, Andrew J. Borst, Robert J. Ragotte, Lukas F. Milles, Basile I. M. Wicky, Nikita Hanikel, Samuel J. Pellock, Alexis Courbet, William Sheffler, Jue Wang, Preetham Venkatesh, Isaac Sappington, Susana Vázquez Torres, Anna Lauko, Valentin De Bortoli, Emile Mathieu, Sergey Ovchinnikov, Regina Barzilay, Tommi S. Jaakkola, Frank DiMaio, Minkyung Baek, and David Baker. De novo design of protein structure and function with RFdiffusion. Nature, 620(7976):1089–1100, July 2023. doi: 10.1038/s41586-023-06415-8. URL 10.1038/s41586-023-06415-8.

[10] Andy Hsien-Wei Yeh, Christoffer Norn, Yakov Kipnis, Doug Tischer, Samuel J. Pellock, Declan Evans, Pengchen Ma, Gyu Rie Lee, Jason Z. Zhang, Ivan Anishchenko, Brian Coventry, Longxing Cao, Justas Dauparas, Samer Halabiya, Michelle DeWitt, Lauren Carter, K. N. Houk, and David Baker. De novo design of luciferases using deep learning. Nature, 614(7949):774–780, February 2023. doi: 10.1038/s41586-023-05696-3. URL 10.1038/s41586-023-05696-3.

[11] Casper A. Goverde, Martin Pacesa, Lars J. Dornfeld, Sandrine Georgeon, Stéphane Rosset, Justas Dauparas, Christian Schellhaas, Simon Kozlov, David Baker, Sergey Ovchinnikov, and Bruno E. Correia. Computational design of soluble analogues of integral membrane protein structures. May 2023. doi: 10.1101/2023.05.09.540044. URL 10.1101/2023.05.09.540044.

[12] Bowen Jing, Stephan Eismann, Patricia Suriana, Raphael John Lamarre Townshend, and Ron Dror. Learning from protein structure with geometric vector perceptrons. In International Conference on Learning Representations, 2021. URL https://openreview.net/forum?id=1YLJDvSx6J4.

[13] Alexander Rives, Joshua Meier, Tom Sercu, Siddharth Goyal, Zeming Lin, Jason Liu, Demi Guo, Myle Ott, C. Lawrence Zitnick, Jerry Ma, and Rob Fergus. Biological structure and function emerge from scaling unsupervised learning to 250 million protein sequences. Proceedings of the National Academy of Sciences, 118(15):e2016239118, 2021. doi: 10.1073/pnas.2016239118. URL https://www.pnas.org/doi/abs/ 10.1073/pnas.2016239118.

[14] Zeming Lin, Halil Akin, Roshan Rao, Brian Hie, Zhongkai Zhu, Wenting Lu, Nikita Smetanin, Robert Verkuil, Ori Kabeli, Yaniv Shmueli, Allan dos Santos Costa, Maryam Fazel-Zarandi, Tom Sercu, Salvatore Candido, and Alexander Rives. Evolutionary-scale prediction of atomic level protein structure with a language model. *bioRxiv*, 2022. doi: 10.1101/2022.07.20.500902. URL https://www.biorxiv.org/content/early/2022/12/21/2022.07.20.500902.

[15] María Sofía Castelli, Paul McGonigle, and Pamela J Hornby. The pharmacology and therapeutic applications of monoclonal antibodies. Pharmacology research & perspectives, 7(6):e00535, 2019.

[16] Rahmad Akbar, Philippe A. Robert, Milena Pavlović, Jeliazko R. Jeliazkov, Igor Snapkov, Andrei Slabodkin, Cédric R. Weber, Lonneke Scheffer, Enkelejda Miho, Ingrid Hobæk Haff, Dag Trygve Tryslew Haug, Fridtjof Lund-Johansen, Yana Safonova, Geir K. Sandve, and Victor Greiff. A compact vocabulary of paratope-epitope interactions enables predictability of antibody-antigen binding. Cell Reports, 34(11):108856, Mar 2021. ISSN 2211-1247. doi: 10.1016/j.celrep.2021.108856. URL 10.1016/j.celrep.2021.108856.

[17] James Dunbar, Konrad Krawczyk, Jinwoo Leem, Terry Baker, Angelika Fuchs, Guy Georges, Jiye Shi, and Charlotte M Deane. Sabdab: the structural antibody database. Nucleic acids research, 42(D1): D1140–D1146, 2014.

[18] Magnus Haraldson Høie, Alissa Hummer, Tobias H. Olsen, Broncio Aguilar-Sanjuan, Morten Nielsen, and Charlotte M. Deane. Antifold: Improved antibody structure-based design using inverse folding, 2024.

[19] Varun R. Shanker, Theodora U. J. Bruun, Brian L. Hie, and Peter S. Kim. Unsupervised evolution of protein and antibody complexes with a structure-informed language model. Science, 385(6704):46–53, 2024. doi: 10.1126/science.adk8946. URL https://www.science.org/doi/abs/10.1126/science.adk8946.

[20] Hyun-Soo Cho, Karen Mason, Kasra X. Ramyar, Ann Marie Stanley, Sandra B. Gabelli, Dan W. Denney, and Daniel J. Leahy. Structure of the extracellular region of HER2 alone and in complex with the herceptin fab. Nature, 421(6924):756–760, February 2003. doi: 10.1038/nature01392. URL 10.1038/nature01392.

[21] Neil Houlsby, Andrei Giurgiu, Stanislaw Jastrzebski, Bruna Morrone, Quentin de Laroussilhe, Andrea Gesmundo, Mona Attariyan, and Sylvain Gelly. Parameter-efficient transfer learning for nlp, 2019.

[22] H. M. Berman. The protein data bank. Nucleic Acids Research, 28(1):235–242, January 2000. doi: 10.1093/nar/28.1.235. URL 10.1093/nar/28.1.235.

[23] Diederik P. Kingma and Jimmy Ba. Adam: A method for stochastic optimization, 2014.

[24] H. B. Mann and D. R. Whitney. On a test of whether one of two random variables is stochastically larger than the other. The Annals of Mathematical Statistics, 18(1):50–60, March 1947. doi: 10.1214/aoms/1177730491. URL 10.1214/aoms/1177730491.

[25] R. A. Fisher. On the interpretation of 2 from contingency tables, and the calculation of p. Journal of the Royal Statistical Society, 85(1):87, January 1922. doi: 10.2307/2340521. URL https://doi.org/10. 2307/2340521.

[26] Brian L. Hie, Varun R. Shanker, Duo Xu, Theodora U. J. Bruun, Payton A. Weidenbacher, Shaogeng Tang, Wesley Wu, John E. Pak, and Peter S. Kim. Efficient evolution of human antibodies from general protein language models. Nature Biotechnology, April 2023. doi: 10.1038/s41587-023-01763-2. URL 10.1038/s41587-023-01763-2.

[27] Ahmed Elnaggar, Michael Heinzinger, Christian Dallago, Ghalia Rehawi, Yu Wang, Llion Jones, Tom Gibbs, Tamas Feher, Christoph Angerer, Martin Steinegger, Debsindhu Bhowmik, and Burkhard Rost. Prottrans: Toward understanding the language of life through self-supervised learning. IEEE Transactions on Pattern Analysis and Machine Intelligence, 44(10):7112–7127, October 2022. ISSN 1939-3539. doi: 10.1109/tpami.2021.3095381. URL 10.1109/TPAMI.2021.3095381.

[28] Martin Steinegger and Johannes Söding. Mmseqs2 enables sensitive protein sequence searching for the analysis of massive data sets. Nature biotechnology, 35(11):1026–1028, 2017.

[29] Arian Rokkum Jamasb, Ramon Viñas Torné, Eric J Ma, Yuanqi Du, Charles Harris, Kexin Huang, Dominic Hall, Pietro Lio, and Tom Leon Blundell. Graphein - a python library for geometric deep learning and network analysis on biomolecular structures and interaction networks. In Alice H. Oh, Alekh Agarwal, Danielle Belgrave, and Kyunghyun Cho, editors, Advances in Neural Information Processing Systems, 2022. URL https://openreview.net/forum?id=9xRZlV6GfOX.

[30] Richard W. Shuai, Jeffrey A. Ruffolo, and Jeffrey J. Gray. Generative language modeling for antibody design. December 2021. doi: 10.1101/2021.12.13.472419. URL 10.1101/2021.12.13.472419.

[31] Shifu Chen, Yanqing Zhou, Yaru Chen, and Jia Gu. fastp: an ultra-fast all-in-one FASTQ preprocessor. Bioinformatics, 34(17):i884–i890, 09 2018. ISSN 1367-4803. doi: 10.1093/bioinformatics/bty560. URL 10.1093/bioinformatics/bty560.

[32] Marcel Martin. Cutadapt removes adapter sequences from high-throughput sequencing reads. EMB-net.journal, 17(1), May 2011.

